# The ratio of Wnt signaling activity to Sox2 transcription factor levels predicts neuromesodermal fate potential

**DOI:** 10.1101/2025.01.16.633481

**Authors:** Robert D. Morabito, David Tatarakis, Ryan Swick, Samantha Stettnisch, Thomas F. Schilling, Julia A. Horsfield, Benjamin L. Martin

## Abstract

Neuromesodermal progenitors (NMPs) are a vertebrate cell type that contribute descendants to both the spinal cord and the mesoderm. The undifferentiated bipotential NMP state is maintained when both Wnt signaling is active and Sox2 is present. We used transgenic reporter lines to live-image both Wnt activity and Sox2 levels in NMPs and observed a unique cellular ratio in NMPs compared to NMP-derived mesoderm or neural tissue. We used this unique signature to identify the previously unknown anatomical position of a progenitor population that gives rise to the midline tissues of the floor plate of the spinal cord and the mesodermal notochord. Thus, quantification of the active Wnt signaling to Sox2 ratio can be used to predict and identify cells with neuromesodermal potential. We also developed the auxin inducible degron 2 system for use in zebrafish to test the temporal role that Sox2 plays during midline formation. We found ectopic Sox2 in the presence of Wnt activity holds cells in the undifferentiated floor plate/notochord progenitor state, and that degradation of the ectopic Sox2 is required for cells to adopt a notochord fate.

## Introduction

During vertebrate embryonic development, the anterior forms first and differentiated tissue is progressively added to the posterior end (Kimelman and Martin, 2012; Martin, 2020). A key population of progenitors that provides new cells to the expanding tissues is the neuromesodermal progenitors (NMPs), which reside in the tailbud structure at the posterior-most end of the developing embryo. NMPs in the posterior wall of the tailbud are highly plastic and continue to make germ-layer fate decisions after gastrulation, giving rise to spinal cord, somites, and endothelium as the embryonic axis extends (Kimelman, 2016; Wymeersch et al., 2021).

NMPs co-express the transcription factors *tbxt* (also called *brachyury*) and *sox2* (Martin and Kimelman, 2012; Wymeersch et al., 2021). Tbxt (in zebrafish there are two paralogs *tbxta* and *tbxtb*, for simplicity we will refer to them as Tbxt) activates expression of canonical Wnt signaling ligands, and Wnt signaling in turn activates *tbxt* expression to form an autoregulatory loop (Martin and Kimelman, 2008, 2009, 2010; Yamaguchi et al., 1999). The NMP fate decision to become neural ectoderm or mesoderm is governed in part by the canonical Wnt signaling pathway and the Sox2 transcription factor. High levels of Wnt signaling repress *sox2* expression and induce mesoderm (Kinney et al., 2020; Martin and Kimelman, 2012). On the other hand, in the absence of Wnt signaling and presence of Sox2, NMPs are induced to form neural ectoderm (Kinney et al., 2020). When both Wnt signaling and Sox2 are present, NMPs maintain an undifferentiated state with potential to form neural or mesodermal tissue (Kinney et al., 2020; Martin and Steventon, 2022). This implies that NMPs employ a unique ratio of canonical Wnt signaling activity to Sox2 expression levels that shifts up or down during mesodermal or neural differentiation, respectively.

The tailbud also contains neuromesodermal progenitors that continuously generate the mesodermal notochord and neural floor plate tissues (Catala et al., 1995; Le Douarin and Halpern, 2000; Martin, 2021; Row et al., 2016; Teillet et al., 1998). These cells, which are also called dorsal midline progenitors, co-express *tbxt* and *sox2* and depend on Wnt activity and Sox2 levels for cell fate specification (Row et al., 2016). Notochord fate depends on active Wnt signaling and the absence of *sox2* expression. When Wnt signaling is inhibited or *sox2* expression maintained, dorsal midline neuromesodermal progenitors form only floor plate (Row et al., 2016). Despite the developmental importance of dorsal midline NMPs, it is unclear precisely where these cells are located within the embryo and how they behave during axis elongation and cell fate specification.

The Sox2 transcription factor plays an important role in maintaining pluripotency in mouse epiblast cells (Avilion et al., 2003) and is sufficient along with a small number of other transcription factors to induce pluripotency in differentiated cells when ectopically expressed (Takahashi et al., 2007; Takahashi and Yamanaka, 2006). At later development stages, Sox2 is required for neuroectoderm specification (Bergsland et al., 2011; Bunina et al., 2020; Thomson et al., 2011; Zhang et al., 2019; Zhang and Cui, 2014). In both the direct differentiation of neuroectoderm from embryonic stem cells and the differentiation of spinal cord from NMPs, Sox2 represses mesodermal inducing genes (Koch et al., 2017; Wang et al., 2012; Zhang and Cui, 2014). Our previous work showed that ectopic activation of *sox2* in midline progenitors prevents notochord induction and induces floor plate formation (Row et al., 2016). However, in this context it is unclear whether Sox2 is acting to maintain the NMP state, similar to its role in embryonic stem cells, or if Sox2 is directly inducing the floor plate fate from NMPs akin to its role during neural specification.

We took a multifaceted approach to identifying and characterizing the dorsal NMP population using high resolution imaging and single cell RNA sequencing (scRNA-seq). We also developed the auxin inducible degron 2 system for use in zebrafish to establish the role of Sox2 in dorsal midline NMPs. We found that the Wnt signaling activity to *sox2* expression level ratio can be used to predict and identify cells with neuromesodermal potential, and that dorsal midline NMPs sit at the posterior-most floor plate position. These cells proliferate and migrate posteriorly, entering the G1 phase of the cell cycle before migrating ventrally to join the notochord. scRNA-seq confirmed the existence of a proliferative bipotential NMP population that gives rise to floor plate and notochord, with dorsal midline NMP cells rapidly downregulating *sox2* as they join the notochord. Transient induction of Sox2 in dorsal midline NMPs using the auxin inducible degron 2 system showed that Sox2 maintains cells in the progenitor state, and cells can join the notochord only after Sox2 is degraded.

## Results

### Live imaging of the Wnt:Sox2 ratio in neuromesodermal progenitors and their descendants

Canonical Wnt signaling and the Sox2 transcription factor play critical roles in both the maintenance of the undifferentiated NMP state and NMP-derived lineage specification. When Sox2 is expressed and Wnt signaling is active, cells maintain the undifferentiated state (Kinney et al., 2020). When Sox2 is expressed in the absence of Wnt signaling, NMPs adopt a neural fate (Kinney et al., 2020; Martin and Kimelman, 2012). On the other hand, when Wnt signaling is active in the absence of Sox2, NMPs become mesoderm (Kinney et al., 2020; Martin and Kimelman, 2012). To directly visualize the relative levels of canonical Wnt signaling and *sox2* in NMPs and their descendants, we used transgenic lines that report on *sox2* expression levels and Wnt signaling activity. To visualize *sox2* expression, we used the *sox2-p2a-sfGFP* line, which contains a *p2a-sfGFP* insertion at the 3’ end of the coding region of the endogenous *sox2* gene (Shin et al., 2014). For canonical Wnt activity visualization, we used the *7xTCF.Siam:NLS-mCherry* line, which expresses a nuclear localized mCherry under the control of 7 multimerized TCF binding sites (Moro et al., 2012). We imaged the tailbud of 16-somite stage embryos containing both the *sox2-p2a-sfGFP* and the *7xTCF.Siam:NLS-mCherry* transgenes and quantified the mean pixel intensity of both reporters in individual cells using spinning disc confocal microscopy. We compared expression levels in the NMPs (cells in the posterior wall of the tailbud) with cells in the posterior spinal cord or posterior presomitic mesoderm (PSM) (Fig. 1A-D’’’). We calculated a ratio of Wnt signaling activity divided by *sox2* reporter activity. We found the highest ratio in the PSM, the lowest in the spinal cord, and an intermediate level in the NMPs, consistent with expectations based on prior Wnt and Sox2 functional modulation (Fig.1 E). To ensure that reporter transgenes are providing dynamic readouts of Wnt and *sox2* levels, we performed time-lapse imaging of transgenic cells transplanted into the tailbud of wild-type host embryos (Fig. 1F-G’, Supplemental Movie 1). Quantification of reporter fluorescence as cells transitioned from NMP to PSM showed that the Wnt:*sox2* reporter ratio increased during mesoderm induction (Fig. 1H). Conversely, cells that transitioned from NMP to neural tissue showed a decrease in the Wnt:*sox2* reporter ratio (Fig. 1H).

**Figure 1.**
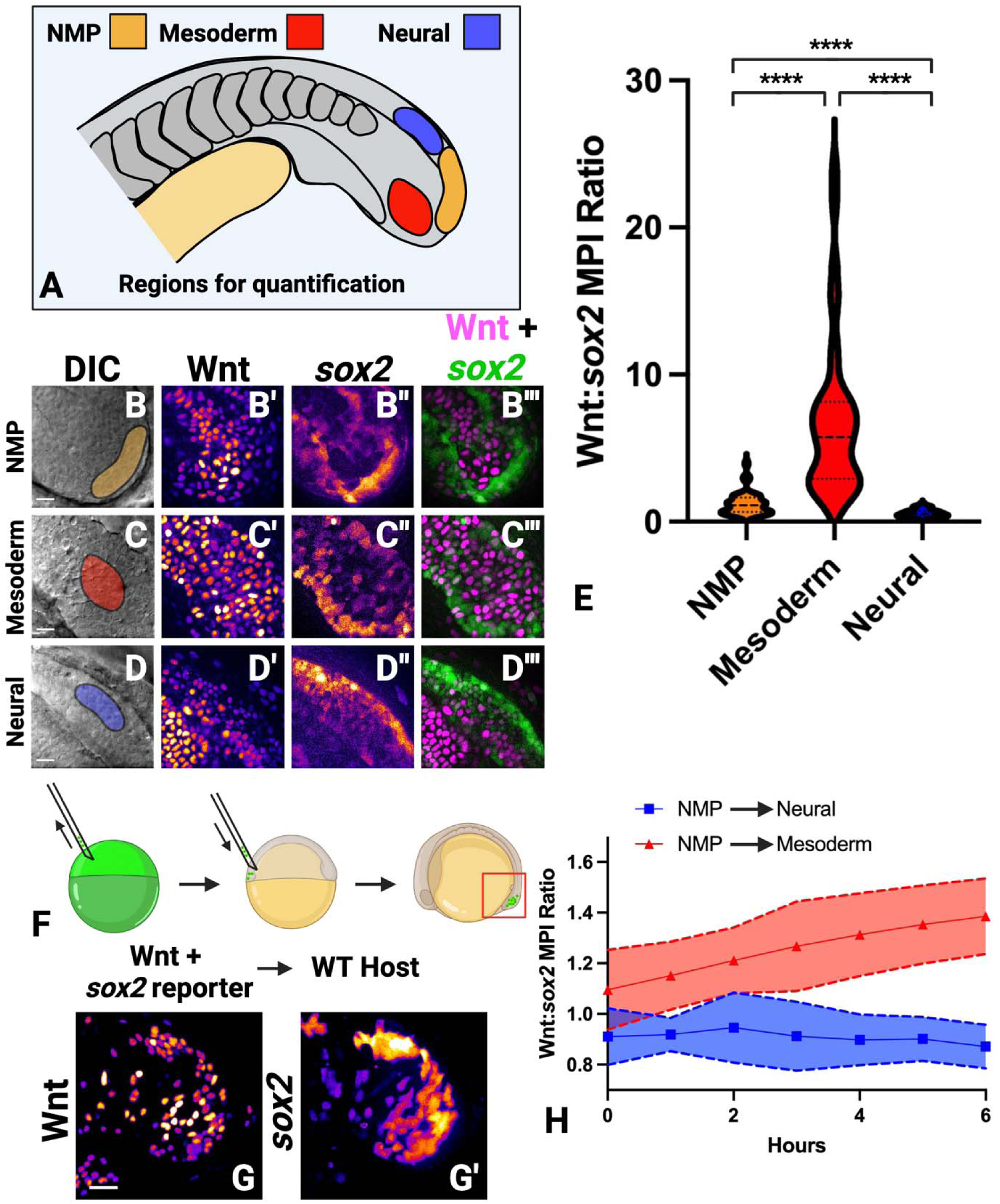
Neuromesodermal progenitors and their descendants have a unique Wnt signaling to Sox2 expression ratio. (A) Embryos containing a single copy of both the *7xTCF-Xla.Siam:nlsmCherry* (Wnt reporter) and *sox2-2a-sfGFP* (*sox2* reporter) were imaged and mean pixel intensities were measured in the NMP, paraxial mesoderm, and neural tube regions of the tailbud (40 NMP, 41 paraxial mesoderm, 30 neural cells measured). (B-D’’’) Representative images of the NMP (B-B’’’), paraxial mesoderm (C-C’’’), and neural tube (D-D’’’) regions are shown. (E) Quantification of the mean pixel intensities were calculated as a ratio of Wnt reporter fluorescence divided by *sox2* reporter fluorescence. The ratio in each region was significantly different than other regions (P<0.0001, unpaired t-test). (F) A cell transplantation method was used to target donor cells with both reporter transgenes into the NMP population of unlabeled wild-type host embryos. (G, G’) Time-lapse imaging was performed, with time-point 0 shown. (H) The mean pixel intensity ratio of Wnt reporter divided by the *sox2* reporter was calculated over time in NMP cells that join the paraxial mesoderm or in NMP cells that join the neural tube, showing dynamic changes in the ratio over time (20 cells measured, 10 that become mesoderm, 10 that become neural). Scale bars = 20μm.

### Identification of a neuromesodermal progenitor-like Wnt:*sox2* ratio in midline progenitors

To determine if the Wnt:*sox2* ratio predicts NMP potential, we quantified the ratio in midline cells in different anterior-posterior or dorsoventral locations with the tailbud. We previously described two midline progenitor populations, one dorsal that gives rise to mesodermal notochord and neural floor plate fates, and a second ventral population that forms either notochord or endodermal hypochord fates (Row et al., 2016). However, it is unclear where exactly these midline progenitors reside relative to one another. By live-imaging of the Wnt:*sox2* ratio in midline cells (Fig. 2A), we found the highest ratio in the notochord and lowest in the hypochord (Fig. 2B-C). On the other hand, cells in the region of the posterior floor plate have a similar Wnt:*sox2* ratio as that of the posterior wall NMPs, suggesting that these cells have potential to become either neural or mesodermal tissue (Fig. 2C, D).

**Figure 2.**
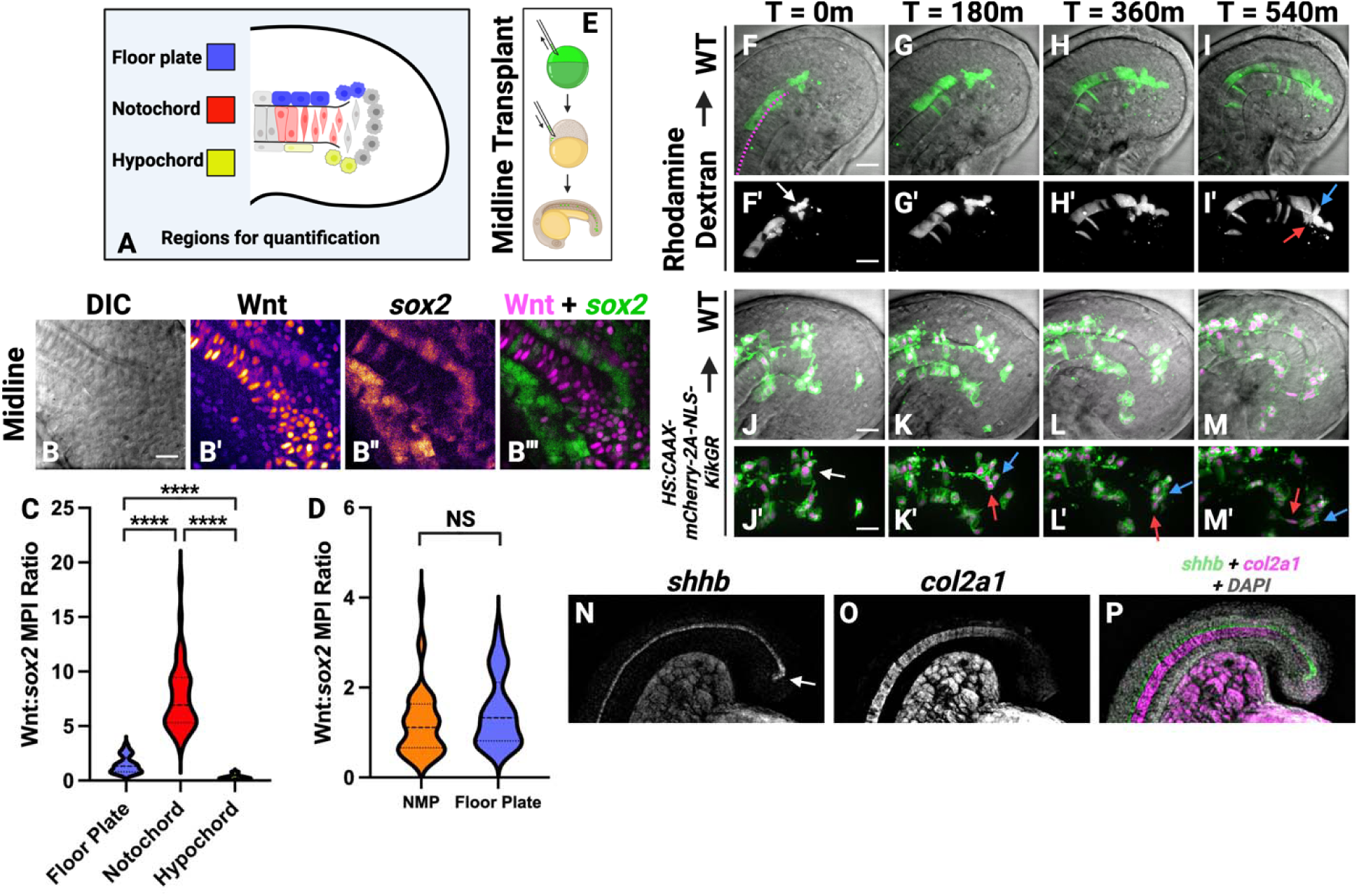
The Wnt:*sox2* ratio identifies dorsal midline progenitors with neuromesodermal potential. (A) Embryos containing a single copy of both the *7xTCF-Xla.Siam:nlsmCherry* (Wnt reporter) and *sox2-2a-sfGFP* (*sox2* reporter) were imaged and mean pixel intensities were measured in the posterior hypochord, notochord, and floor plate regions of the tailbud (42 notochord, 32 floor plate, 28 hypochord cells measured). (B-B’’’) Representative images of the midline tissues are shown. (C) Quantification of the mean pixel intensities were calculated as a ratio of Wnt reporter fluorescence divided by *sox2* reporter fluorescence. The ratio in each region was significantly different than other regions (P<0.0001, unpaired t-test). (D) The ratio between the floor plate and the NMPs was not statistically different. (E) A cell transplantation method was used to target labeled donor cells to the midline progenitors of unlabeled wild-type host embryos. (F-I’) Wild-type cells labeled with rhodamine dextran were transplanted into the midline progenitor population of unlabeled wild-type host embryos and time-lapse imaging was performed beginning at the 16-somite stage. (F) The border between the notochord and floor plate is indicated by a pink dashed line. Cells continuous with the posterior floor plate (white arrow, F’) move posteriorly relative to the notochord. Cells within this population contribute to both floor plate (I’, blue arrow) and notochord (I’, red arrow), with cells moving ventrally when forming the notochord (N=6 movies). (J-M’) Transgenic *HS:CAAX-mCherry-2a-NLS-KikGR* cells were transplanted into the midline progenitor domain of unlabeled wild-type host embryos and transgene expression was induced at bud stage. Time-lapse movies were made beginning at the 16-somite stage (J-M’, membranes are labeled in green and nuclei in magenta). Cells continuous with the posterior floor plate (white arrow, J’) move posteriorly relative to the notochord. Cells within this population contribute to floor plate and notochord (K-M’, red arrows show a notochord forming cell and blue arrows point to floor plate forming cells, N=4 movies). (N-P) Hybridization chain reaction was performed on embryos at the 16-somite stage using probes for the floor plate marker *shhb* and the notochord marker *col2a1*. At the posterior end of the floor plate the expression domain of *shhb* turns sharply into where the notochord progenitors reside (N, white arrow). Scale bars = 20μm.

### The posterior-most region of the floor plate is a neuromesodermal progenitor population that gives rise to the definitive floor plate and notochord

To test if cells in the posterior-most floor plate have the potential to give rise to both neural and mesodermal cells, we performed midline-directed transplants, which target cells specifically to tailbud midline-progenitor cells (Fig. 2E) (Halpern et al., 1995; Row et al., 2016). We first transplanted cells from rhodamine dextran-injected donor embryos into unlabeled wild-type host embryos (Fig. 2F-I’, Supplemental Movie 2). When transplanted cells were located in the position of presumptive floor plate progenitors (i.e. more dorsal in the tailbud), they migrated posteriorly with respect to grafted cells in the notochord. Cells in this posterior-most putative floor plate (Fig. 2F’, white arrow) were then either left behind the rapidly migrating posterior population and adopted a bonafide floor plate fate or migrated posteriorly and ventrally to join the notochord (Fig. 2I’, blue arrow indicates floor plate cell, red arrow indicates notochord cells). We also performed transplants using the *HS:CAAX-mCherry-2a-NLS-KikGR* transgenic line, which labels the plasma membrane with mCherry and the nucleus with KikGR (Goto et al., 2017), as donor embryos (Fig. 2J-M’, Supplemental Movie 3). We saw the same results, with cells in the posterior-most region of the ventral midline ectoderm (Fig. 2J’, white arrow) becoming either floor plate or notochord (Fig. 2J-M’, blue arrow indicates floor plate, red arrow indicates notochord). To provide further evidence that the posterior floor plate-like progenitors can generate notochord cells, we performed hybridization chain reaction (HCR) to detect the spatial localization of the mRNA for the classic floor plate marker *shhb* and the notochord specific marker *col2a1* (Ekker et al., 1995; Yan et al., 1995). We observed that the entirety of the floor plate expresses *shhb* with the posterior-most expression domain extending ventrally into cells that appear to cap the posterior notochord domain (Fig. 2N-P, white arrow). This, along with the time-lapse confocal imaging of transplanted cells, suggests that cells expressing *shhb* become incorporated into the notochord, after which they turn off *shhb* and activate notochord specific gene expression.

### Single-cell RNA-seq reveals the cellular landscape of the tailbud and identifies midline progenitor cells

To comprehensively profile the many cell types and progenitor cell states in the developing tailbud, we generated a single-cell transcriptomic atlas of the zebrafish tailbud. 30 Tailbuds were isolated from 16 hpf embryos by manual dissection and then dissociated into single cells by trypsin and collagenase P digestion. Cell suspensions were pooled prior to generation of libraries. Single-cell libraries were constructed on the 10X Chromium platform and sequenced on an Illumina HiSeq2500 (Fig. 3A). Following filtering for low quality cells, 11,138 total single-cell transcriptomes were obtained with an average of about 3500 genes detected per cell.

**Figure 3.**
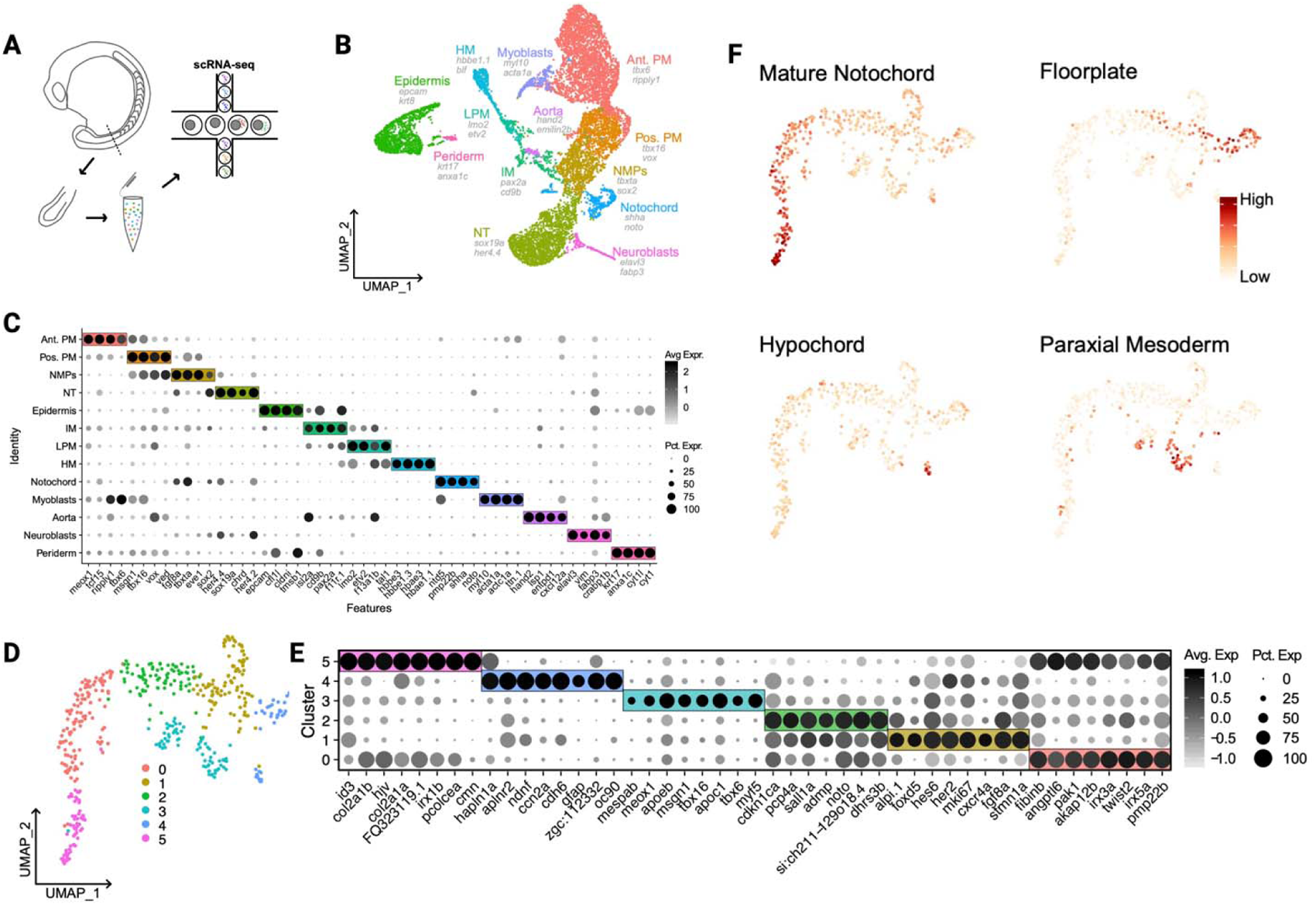
scRNA-seq of the developing vertebrate tailbud identifies neuromesodermal progenitors and their derivatives. (A) Schematic showing procedure for tailbud single cell isolation and sequencing. (B) UMAP embedding of 11,138 single cells sequenced from the tailbud. Cell types are labeled, and two markers for each cell type are included below labels. (C) Dotplot showing top 4 marker genes in each cell type by differential expression analysis. Colored boxes correspond to cell type. Greyscale indicates average expression and dot size indicates percent of expressing cells within each cell type. Marker genes were determined by Wilcoxon rank sum test. (D) UMAP embedding of all cells assigned a midline tissue identity in the full tailbud analysis. Unbiased clustering resulted in 5 distinct clusters. (E) Dotplot showing top 4 marker genes in each cell cluster shown in C by differential expression analysis. Colored boxes correspond to cell type. Greyscale indicates average expression and dot size indicates percent of expressing cells within each cell type. (F) UMAP overlay of gene scoring for cell-type specific markers.

Following regression to remove cell cycle effects, dimensionality reduction and clustering revealed 13 major cell types based on their expression of canonical marker genes (Fig. 3B-C). Epidermal cells marked by expression of *epcam* and *krt8*, showed distinct signatures from peridermal cells that express *krt17* and *anxa1c*. Neural tube (NT) cells expressed *sox19a* and multiple *her* genes, with more mature neuroblasts expressing *elavl3*. Notochord cells were easily identified by expression of *noto* and *shha*. Two distinct populations of paraxial mesoderm (PM), which we dubbed “anterior” (more mature) and “posterior” (less mature) shared high expression of *msgn1* and *tbx16*, but the posterior PM had much higher expression of *ripply1*, and the posterior PM expressed high levels of *vox* and *ved*. A distinct cluster of NMPs sits sandwiched between the NT and PM populations in UMAP space, an observation similar to other zebrafish scRNA-seq studies (Genuth et al., 2023; Lange et al., 2024).

To try and identify midline progenitor cells that give rise to notochord, hypochord, and floor plate, we further subclustered the notochord cluster. Following dimensionality reduction, we found 6 clusters of cells with distinct expression profiles (Fig. 3D-E). By computing gene scores (Tirosh et al., 2016) across the data set for canonical notochord, floor plate, and hypochord marker genes (Fig. 3F). Clusters 0 and 4 showed enrichment for markers of notochord, while parts of clusters 1 and 4 were enriched for floor plate markers. A small population assigned to cluster 4 was enriched for a hypochord signature. Cluster 3 showed enrichment for PM, suggesting these cells may be escaping Noto mediated repression of PM specific gene expression (Amacher and Kimmel, 1998). These results confirm that the tailbud contains all three midline progenitor-derived tissues.

To better understand the connections between these midline progenitor-derived cell types, we next performed trajectory and pseudotime analyses. Dimensionality reduction and trajectory inference in Monocle3 predicted the expected connections between clusters (Fig. 4A-B). The trajectory also implied a branching point between floor plate and notochord fates originating from cluster 3, so pseudotime was calculated with this cluster as the starting node (Fig. 4C). Floor plate marker *sox2* increased along the trajectory branch leading to cluster 4, while in the trajectory leading to cluster 5, expression of this gene drops off rapidly (Fig 4D). On the other hand, the notochord marker *tbxta* was maintained along the trajectory branch leading to cluster 5 and decreased along the trajectory to cluster 4 (Fig. 4D). Cells in cluster 1 showed co-expression of *sox2* and *tbxta*, consistent with this being the midline NMP population (Fig. 4E). We identified *fgf8a* as one of the most enriched genes expressed in NMPs in the whole tailbud (Fig. 3). *fgf8a* showed specific expression in the putative midline NMP cells of cluster 1, and overlapped with *sox2* in this region (Fig. 4F). The transcription factor *noto* was expressed in midline and notochord progenitors - loss of Noto function results in reduced notochord and floor plate (Talbot et al., 1995). We found strong overlap of *fgf8a* and *noto* in cluster 1, identifying a potential novel signature of the midline NMPs (Fig. 4G). To provide further evidence that cluster 1 represents midline progenitors, we performed RNA velocity analysis which uses high-dimensional vectors based on inferred transitional cell states to predict future cell fates (La Manno et al., 2018). This suggests cluster 1 as an origin for both notochord and floor plate (Fig. 4H).

**Figure 4.**
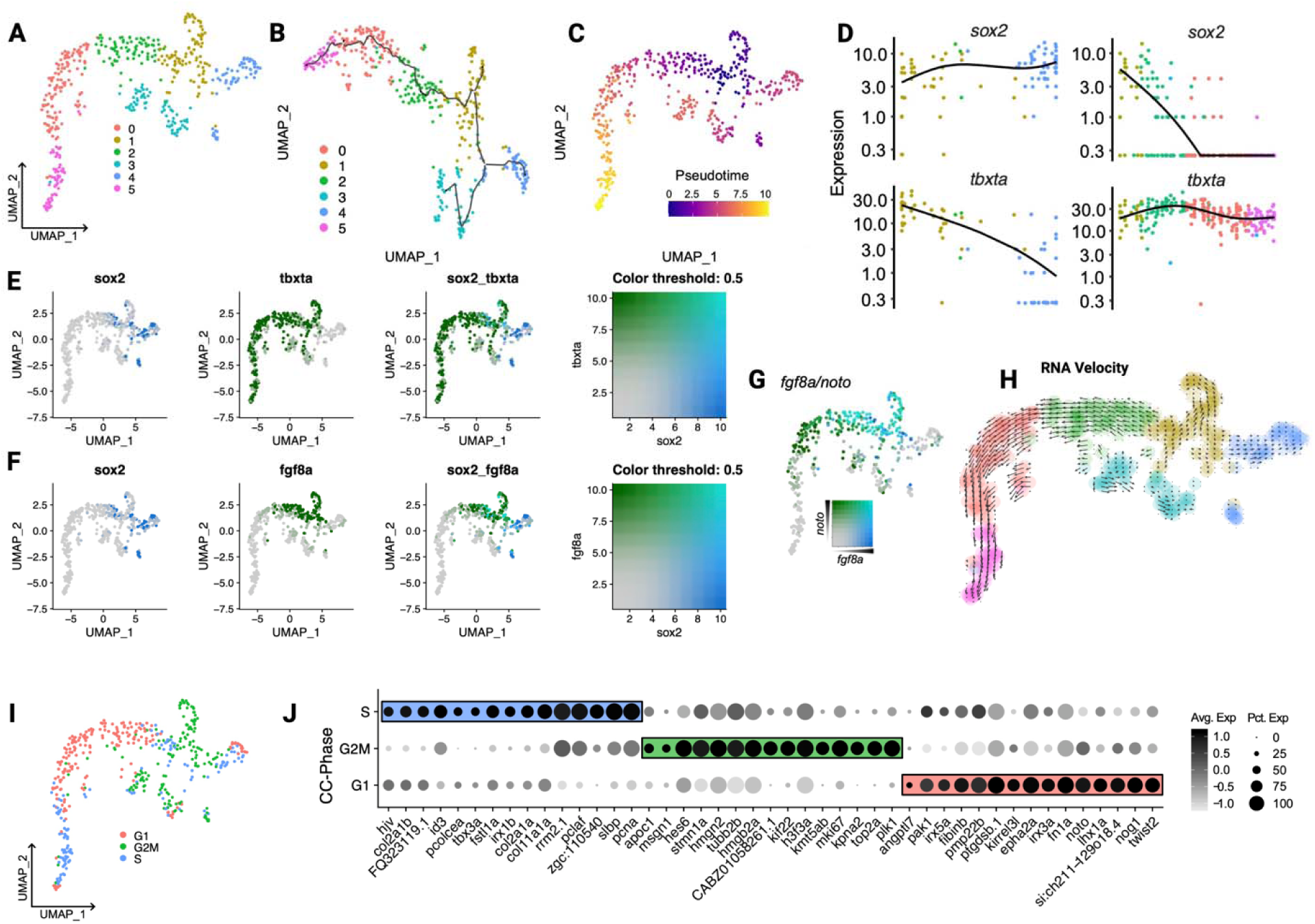
Pseudotime and RNA velocity analysis details emergence of notochord and floor plate and identifies a molecular signature of midline progenitor cells. (A) The UMAP embedding of all cells assigned a midline tissue identity is shown again as a reference. (B) UMAP embedding and trajectory computed in Monocle3. (C) Pseudotime values computed from differentiation trajectory with putative MPC population (cluster 1) as the starting node overlayed with Seurat UMAP. (D) Local regression for NMP markers *sox2* and *tbxta* across the floor plate trajectory branch (cluster 1 to cluster 4) and notochord trajectory branch (cluster 1 to cluster 5). (E) UMAP overlayed with expression for both of the NMP markers *sox2* and *tbxta* or (F) the NMP marker *fgf8a* and *sox2*. Overlapping expression is observed in cluster 1. (G) UMAP overlayed with expression of *fgf8a* and the notochord and midline progenitor marker *noto*. (H) RNA velocity analysis identifies cluster 1 as the putative MPC population. (I) UMAP colored by cell cycle states determined by cell cycle scoring. G1, S, and G2/M stages are not evenly distributed among clusters. (J) Dotplot showing top 15 marker genes in each cell-cycle state by differential expression analysis. Colored boxes correspond to cell type. Greyscale indicates average expression and dot size indicates percent of expressing cells within each cell type.

We analyzed the cell cycle phases of individual cells and colored the UMAP according to each phase (Fig. 4I). G1, S, G2, and M phases of the cell cycle were not evenly distributed amongst cells, but enriched in distinct clusters. For example, the forming and recently formed notochord progenitors in clusters 2 and 0 were almost exclusively in the G1 phase, and more mature notochord cells were predominantly in S phase. This is consistent with previous work using the FUCCI live cell cycle transgenes to show synchronization of the posterior notochord in G1 phase followed by entry of cells into S phase as they mature (Sugiyama et al., 2014; Sugiyama et al., 2009). The cells in clusters 1 and 4 representing midline progenitors and floor plate, respectively, were mostly in the S and G2/M phases of the cell cycle, indicating that these cells actively cycle. Collectively the midline cells in G1 phase expressed genes associated with an immature notochord or notochord progenitor state included *twist2* and *lhx1a* (Fig. 4J). These expression profiles suggest the cell cycle phases may be indicative of differential acquisition of a notochord or floor plate identity (Fig. S1).

### Midline neuromesodermal progenitors are proliferative and enter G1 as they form notochord

The scRNA-seq analyses indicated that different midline cell populations have distinct cell cycle phase characteristics (Figs. 4, S1). Previous analyses of the cell cycle states of NMPs in zebrafish embryos suggested that notochord progenitors arrest in the G1 phase of the cell cycle (Adikes et al., 2020; Sugiyama et al., 2014; Sugiyama et al., 2009). To determine cell cycle states of midline NMPs in live zebrafish embryos, we used a ratiometric Cyclin dependent kinase (Cdk) sensor transgenic line that we previously generated (Adikes et al., 2020; Morabito et al., 2021). This sensor consists of a fragment of human DNA helicase B (DHB) fused to mNeonGreen (mNG). In the absence of Cdk activity, DHB localizes to the nucleus. Phosphorylation of DHB by Cdk induces nuclear export, such that the sensor shifts from nuclear localization to cytoplasmic localization as the cell cycle progresses and Cdk activity increases (Spencer et al., 2013). The transgene also contains a histone H2B fused to mScarlet, which is separated from the DHB by the viral 2A peptide. The transgene is under the control of the *hsp70l* heat-shock inducible promoter for strong ubiquitous expression (Fig. 5A).

**Figure 5.**
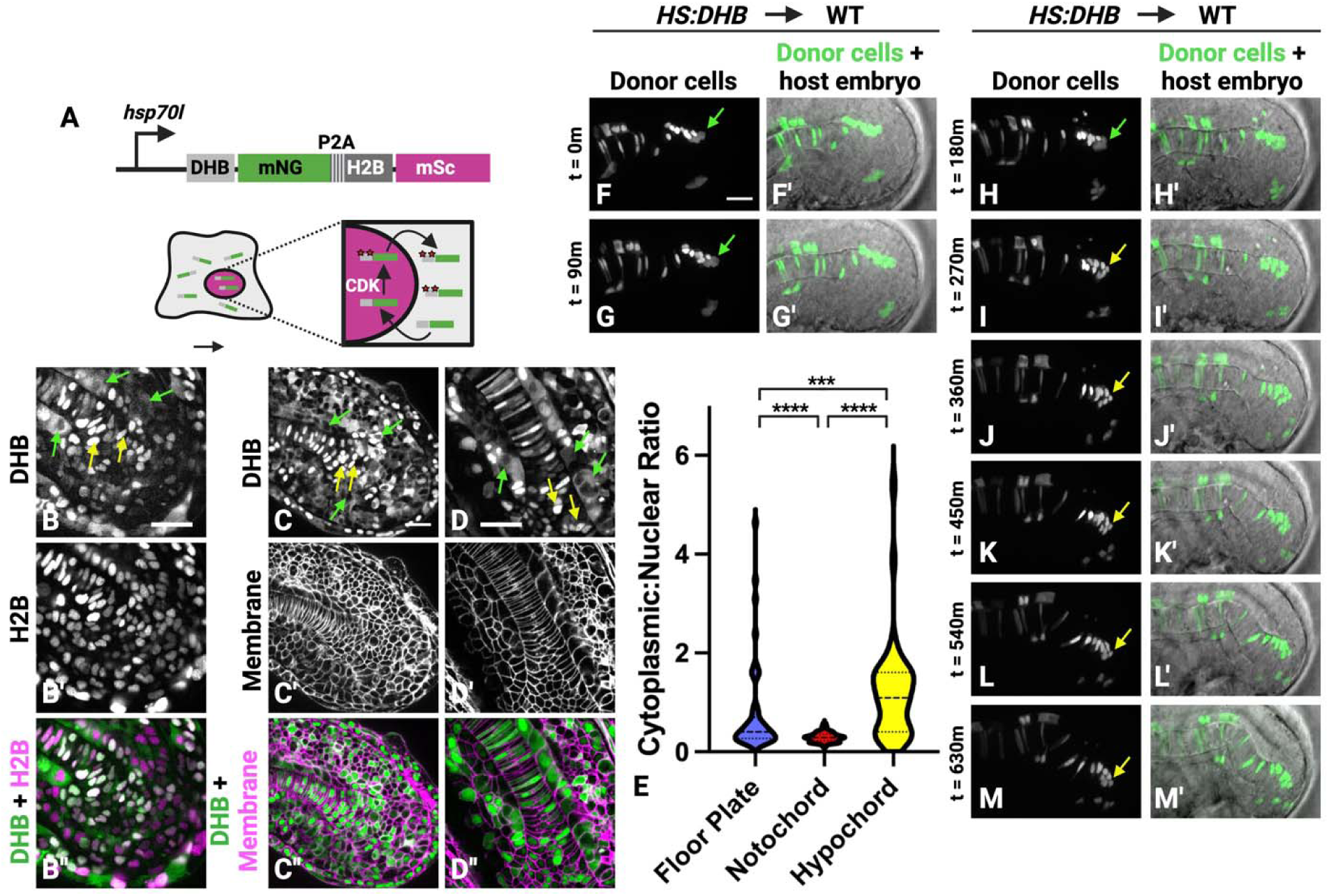
The dorsal midline NMPs are a proliferative cell population. (A) A schematic shows the Cdk sensor transgenic construct used to assess cell cycle state in live cells. (B-B’’’) The *HS:DHB-mNG-p2a-H2B-mSc* transgenic line at the 26-somite stage shows localization of the DHB Cdk sensor (B), nuclei (B’), and are overlayed in B’’. Yellow arrows point to examples of G1 phase cells with nuclear DHB localization, and green arrows point to G2 phase cells with cytoplasmic DHB localization. (C-D’’) The *HS:DHB-mSc-p2a-H2B-miRFP670* was crossed to the plasma membrane reporter line *ubb:lck-mNG* and imaged at the 22-somite stage (C-C’’) and at 24 hpf (D-D’’). Again, yellow arrows point to examples of G1 phase cells with nuclear DHB localization, and green arrows point to G2 phase cells with cytoplasmic DHB localization. (E) Quantification of the cytoplasm:nuclear ratio of DHB at the 26-somite stage shows that notochord cells are in G1 while posterior floor plate and hypochord cells are cycling (118 notochord, 52 floor plate, 43 hypochord cells measured). (F-M’) Midline progenitor targeted transplantation and time-lapse imaging using *HS:DHB-mSc-p2a-H2B-miRFP670* donor cells show that the dorsal midline NMPs are in various phases of the cell cycle but arrest in G1 before entering the notochord domain (F-M’). Green arrows show location of proliferative cells at the dorsal midline NMP migration front (F-H’), which divide and enter G1 (yellow arrows) followed by ventral migration into the notochord domain (J-M’). Scale bars = 20μm.

The cytoplasmic to nuclear ratio of DHB was determined for cells occupying the putative posterior floor plate, notochord, and hypochord in the tailbud (Fig. 5B-E). As previously reported, posterior notochord progenitor cells were held in the G1 phase of the cell cycle (Fig. 5B, C, D, yellow arrows, E). In contrast, floor plate and hypochord progenitors had significantly higher cytoplasmic to nuclear DHB ratio, indicating that these populations proliferate (Fig. 5B, C, D, green arrows, E). To observe cell cycle dynamics during the transition from midline NMP to notochord, we performed transplants targeting DHB transgenic cells to the midline of wild-type host embryos and made time-lapse movies (Fig. 5F-M’, Supplemental Movie 4). The midline NMPs (in the posterior-most region of the floorplate) actively cycle (Fig. 5F-H, green arrows). Cells that join the notochord do so after cell division when they are in the G1 phase, indicating that they are in G1 before undergoing convergence and extension movements (Fig. 5I-M, yellow arrows).

### Lowering Wnt signaling or sustaining sox2 expression maintains cells in the midline neuromesodermal progenitor population

We previously showed that within the dorsal midline progenitor population, the induction of notochord fate requires active Wnt signaling as well as repression of *sox2* expression (Row et al., 2016). Inhibiting Wnt signaling or activating *sox2* expression using heat-shock inducible transgenes at 10 hpf (bud stage) causes dorsal midline progenitors to shift to give rise predominantly to floor plate (Row et al., 2016). In order to confirm that this occurs at later stages of axis extension, we repeated these experiments by transplanting cells from heat-shock inducible *sox2* transgenic embryos (*HS:sox2*) into the midline progenitors of wild-type host embryos, heat-shock induced *sox2* expression at the bud stage, and imaged embryos at 24 hpf.

In controls, in which we transplanted wild-type cells to wild-type host embryos, donor cells contributed to both the floor plate and notochord (Fig. 6A-A’’’). However, in contrast, *HS:sox2* donor cells only contributed to floor plate and not notochord (Fig. 6B-B’’’). In order to examine the behavior of dorsal midline progenitors over time under these conditions, we transplanted cells from heat-shock inducible *tcf*Δ*C* (*HS:tcf*Δ*C*) transgenics, which express a dominant negative inhibitor of the canonical Wnt signaling pathway (Martin and Kimelman, 2012), or *HS:sox2* transgenic cells into wild-type host embryos, targeting them to the midline progenitor population. Host embryos were heat-shocked at bud stage and time-lapse movies were made beginning at the 12-somite stage. Again, control experiments where wild-type cells were transplanted to wild-type host embryos showed cells in the dorsal midline NMPs contributing to both notochord and floor plate (Fig. 6C-F’, Supplemental Movie 5). In contrast, transplanted cells in which *sox2* expression was activated (Fig. 6G-J’, Supplemental Movie 6) or Wnt signaling was inhibited (Fig. 6K-N’, Supplemental Movie 7) remained at the migratory front of the dorsal midline NMPs but failed to move ventrally and join the notochord (white arrows). Over time these cells contributed only to the floor plate.

**Figure 6.**
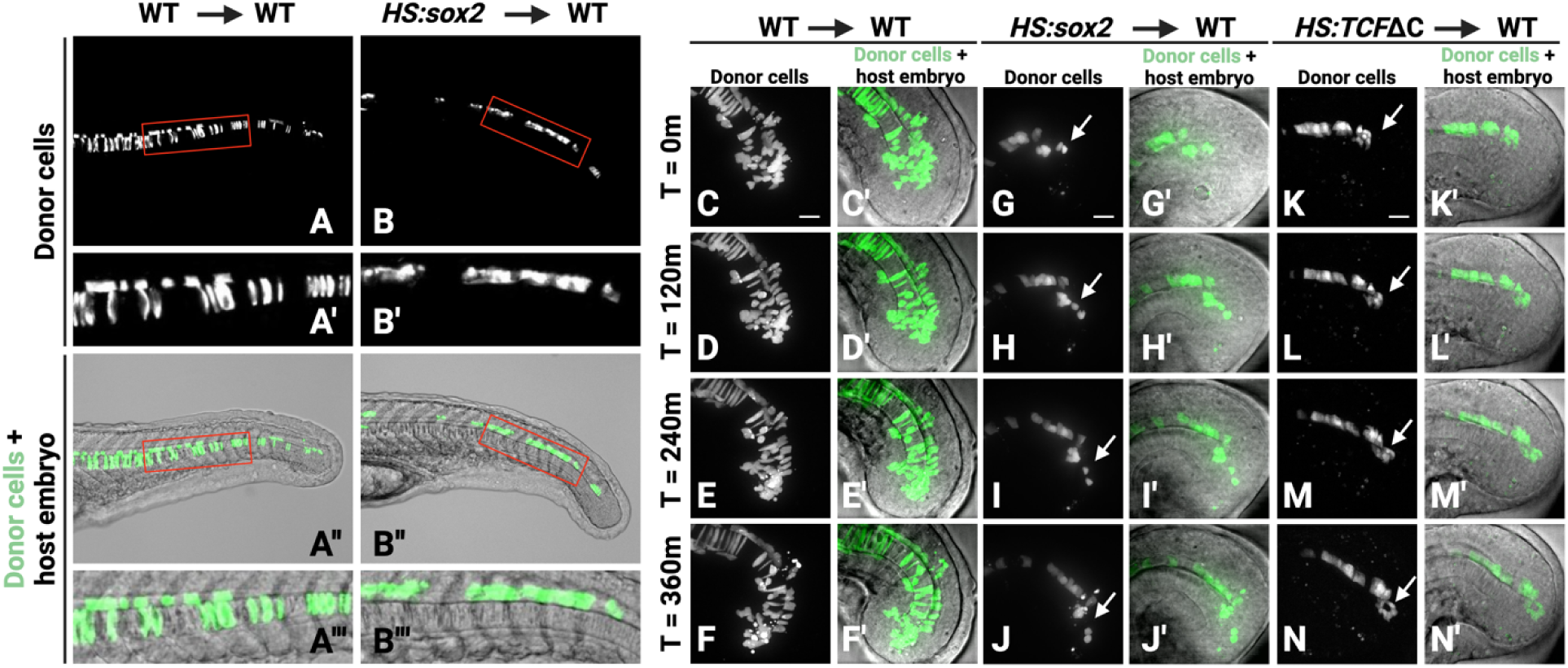
Changing the Wnt:*sox2* ratio prevents the dorsal midline NMP to notochord transition. (A-A’’’) Transplanting wild-type rhodamine dextran labeled cells into the midline progenitor region of host embryos shows transplanted cells contributing to both the floor plate and notochord (red boxes in A and A’’ show regions in A’ and A’’’, respectively N=7). (B-B’’’) When the *sox2* level is elevated in transplanted cells, using the *HS:sox2* transgenic line, cells stay in the dorsal midline progenitor zone but fail to generate notochord (red boxes in B and B’’ show regions in B’ and B’’’, respectively, N=11). Embryos were imaged at 24 hpf and *sox2* was induced at bud stage. (C-F’) Time-lapse imaging of wild-type cells transplanted into the midline progenitor region of wild-type host embryos shows cells in the dorsal midline NMP region contributing to floor plate and notochord. (G-N’) However, when transgenic cells are transplanted into wild-type host embryos and *sox2* is activated (using the *HS:sox2* line) (G-J’) or Wnt signaling is inhibited (using the *HS:tcf*Δ*C* line) (K-N’) at bud stage, cells remain at the migratory front of the dorsal midline NMP region but do not join the notochord (white arrows). Time-lapse imaging was started at the 12-somite stage and continued for 360 minutes. Scale bars = 20μm.

### Developing the auxin inducible degron 2 system in zebrafish

The scRNA-seq analyses show that *sox2* expression is rapidly downregulated as midline progenitors are specified as notochord in the tailbud midline. To confirm this we performed HCR using probes for *sox2* and the notochord marker *col2a1* and observed *sox2* expression in cells surrounding the notochord but not in notochord cells (Fig. 7A-C). Inducing *sox2* expression using the *HS:sox2* transgene caused dorsal midline progenitors to give rise to floor plate (or hypochord) but not notochord (Fig. 6). Sox2 could directly induce floor plate fate, or alternatively Sox2 activity might maintain the undifferentiated NMP state and causing cells to adopt a floor plate fate by default when they cannot join the notochord. We observed in a minority of HS:sox2 midline directed transplants that cells would resume forming notochord at the very posterior tip, presumably when the induced Sox2 turned over, suggesting that Sox2 is maintining the undifferentiated NMP state (Fig. S2). To further distinguish between these two models, we induced *sox2* expression with a heat-shock inducible transgene, the protein produced by which can then be rapidly degraded. If Sox2 directly induces floor plate fate, cells in which Sox2 has been induced and then degraded should still fail to join the notochord. However, if Sox2 maintains the undifferentiated NMP state, then degradation of the induced Sox2 should allow cells to resume notochord contribution.

**Figure 7.**
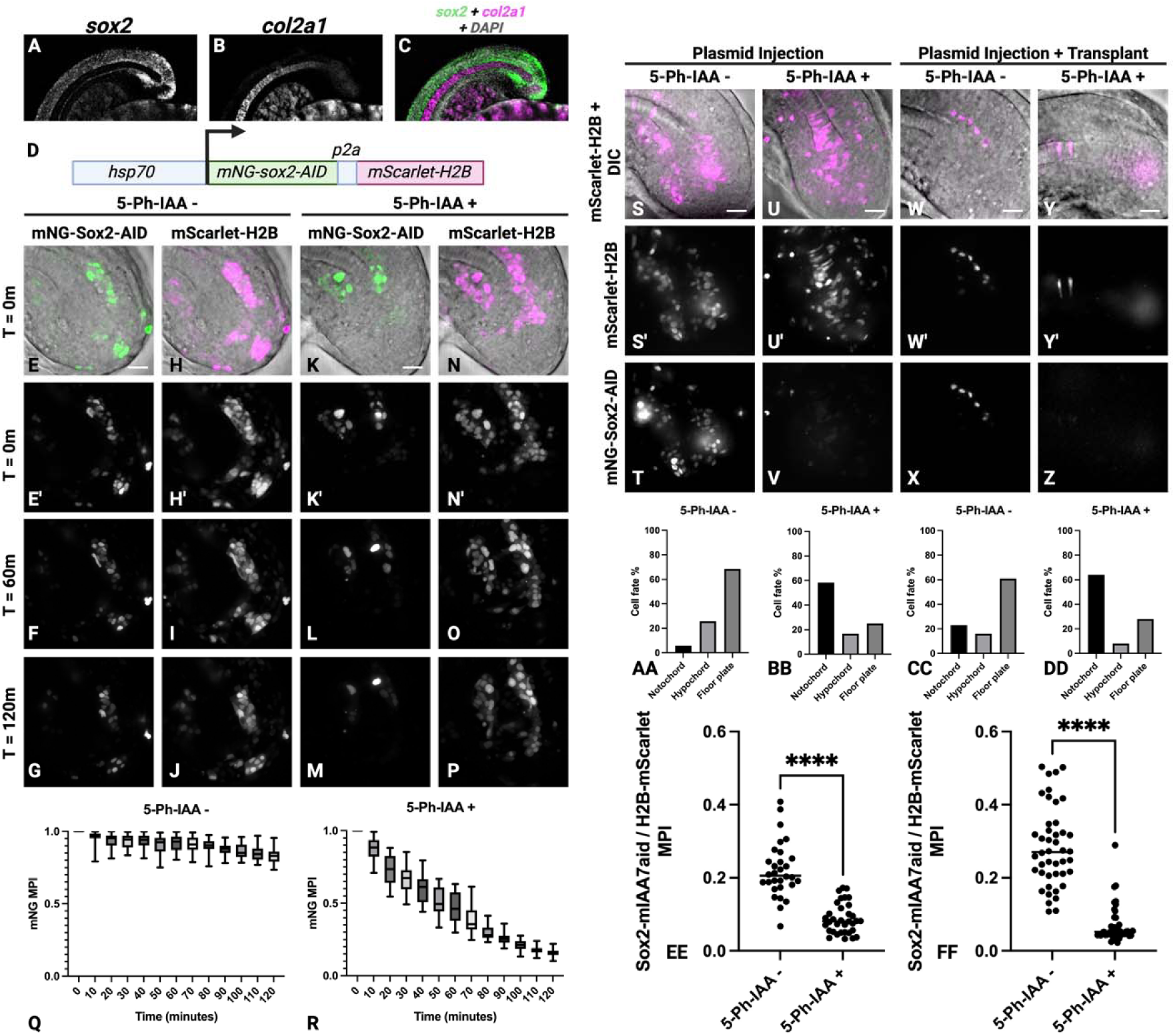
Sox2 maintains dorsal midline NMPs in a bipotential progenitor state. (A-C) Hybridization chain reaction was performed on embryos at the 16-somite stage using probes for *sox2* and the notochord marker *col2a1*. (D) The plasmid construct used to temporally induce expression of *mNG-sox2-AID* and *mScarlet-H2B*. (E-J) Plasmid injected embryos were heat-shocked at shield stage and timelapse imaged for 2 hours. In the absence of 5-Ph-IAA, mNG-Sox2-AID levels are maintained (E-G), as are mSc-H2B levels (H-J). (Q) Mean pixel intensity of mNG-Sox2-AID in the absence of 5-Ph-IAA are plotted at 10-minute intervals (N=30 cells). (K-P) In the presence of 5-Ph-IAA, mNG-Sox2-AID levels are rapidly depleted immediately after 5-Ph-IAA addition (K-M), whereas mScarlet-H2B levels are maintained (N-P). (R) Mean pixel intensities of mNG-Sox2-AID in the presence of 5-Ph-IAA are plotted at 10-minute intervals (N=31 cells). (S-V) Plasmid injected embryos were heat-shocked at shield stage and treated with or without 5-Ph-IAA beginning at bud stage. Embryos were imaged at the 16-somite stage. (S-T, AA, EE) In the absence of 5-Ph-IAA, mNG-Sox2-AID levels are maintained (EE, N=30 cells) and cells join the floor plate and hypochord but not the notochord (S-T, AA, N=35 cells). (U-V, BB, EE) In the presence of 5-Ph-IAA, mNG-Sox2-AID is depleted (EE, N=34 cells) and notochord contribution resumes (U-V, BB, N=36 cells, P<0.0001 chi square test). (W-Z) Plasmid injected embryos were used as donor embryos in midline directed transplants, and host embryos were heat-shocked at the shield stage and treated with or without 5-Ph-IAA beginning at bud stage. Embryos were imaged at the 16-somite stage. (W-X, CC, FF) In the absence of 5-Ph-IAA, mNG-Sox2-AID levels are maintained (FF, N=44 cells) and cells join the floor plate and hypochord but not the notochord (W-X, CC, N=36 cells). (Y-Z, DD, FF) In the presence of 5-Ph-IAA, mNG-sox2-AID is depleted (FF, N=45 cells) and notochord contribution resumes (Y-Z, DD, N=41 cells, P<0.0001 chi square test). Scale bars = 20μm.

To perform this experiment, we adopted the auxin-inducible degradation 2 system (AID2) for use in zebrafish (Yesbolatova et al., 2020). The original AID system utilizes the plant specific f-box protein Tir1, which interacts with the auxin inducible degron (AID) only in the presence of the plant hormone auxin (Nishimura et al., 2009). AID-tagged proteins in the presence of Tir1 can be degraded rapidly upon addition of auxin. However, drawbacks of the AID system include: 1) leaky degradation of AID tagged proteins, and 2) requirements for high auxin concentrations. The AID2 system uses a mutant Tir1 protein and the auxin derivative 5-Ph-IAA, which together prevent leaky degradation and work efficiently at much (670x) lower ligand concentrations (Yesbolatova et al., 2020). When BFP tagged Tir1 mRNA was co-injected with an AID tagged mNG fluorescent protein mRNA, fluorescence in the embryo was rapidly lost upon 5-Ph-IAA addition (Fig. S3). This degradation was more efficient when the AID was replaced with the mIAA7 degron (Fig. S3), which has been shown in mammalian cells and C. elegans to increase degradation efficiency (Li et al., 2019; Sepers et al., 2022).

We next generated a heat-shock inducible Tir1-BFP transgenic line and observed similar degradation kinetics when AID or mIAA7 tagged mNG mRNAs were injected and embryos were treated with or without 5-Ph-IAA (Fig. S4). To modulate Sox2 activity, we designed a heat-shock inducible plasmid to drive a *mNG-sox2* fusion tagged with the mIAA7 degron. This construct also has a 2A peptide that separates a mScarlet-H2B fusion, which allows for tracking of cells that expressed mNG-Sox2 but may no longer exhibit mNG fluorescence (Fig. 7D). This *sox2* plasmid was co-injected with a heat-shock inducible Tir1-mScarlet fusion plasmid. Plasmids are inherited mosaically when injected into 1-cell stage zebrafish embryos. Injected embryos were heat-shocked at the shield stage and then treated with 5-Ph-IAA at the 8-12 somite stage and cells tracked with time-lapse imaging (Fig. 7E-P). Like degron tagged mNG alone, the degron tagged mNG-Sox2 was degraded rapidly in the presence of 5-Ph-IAA, whereas mScarlet fluorescence was maintained (Fig. 7E-R).

### Sox2 maintains the undifferentiated midline NMP state

To determine if exogenous Sox2 directly induces floor plate fate in dorsal midline progenitors or alternatively maintains an undifferentiated progenitor fate, we injected the heat-shock inducible *mNG-sox2-AID* construct along with *HS:tir1-mScarlet* and heat-shocked injected embryos at shield stage. Embryos were then treated with 5-Ph-IAA (or DMSO vehicle in controls) at bud stage and imaged at the 18-somite stage (Fig. 7S-V). In vehicle treated embryos there was very little contribution of plasmid containing cells to the notochord, and instead they remained in either the floor plate or hypochord (Fig. 7S-T, AA). On the other hand, depletion of the induced Sox2 allowed plasmid containing cells to contribute to the notochord (Fig. 7U-V, BB). This was accompanied by a significant loss of mNG-Sox2 fluorescence relative to the mScarlet-H2B fluorescence in individual plasmid containing cells (Fig. 7EE). As a complementary approach, we also performed a similar set of experiments, but in this case used plasmid-injected embryos as donors for midline directed transplants into wild-type host embryos. We observed similar results, with low notochord contribution of plasmid containing transplanted cells in the absence of 5-Ph-IAA, and increased contribution when 5-Ph-IAA is added (Fig. 7W-Z, CC, DD). This was again accompanied by a significant loss of mNG-Sox2 fluorescence relative to the H2B-mScarlet fluorescence in individual plasmid containing cells (Fig. 7FF). These results indicate that Sox2 does not directly induce floor plate fate in dorsal midline progenitors, but rather maintains the undifferentiated progenitor state. The results also add further evidence that *sox2* must be repressed before midline progenitor cells can become notochord.

## Discussion

### An updated model for midline tissue formation from midline progenitor cells

Our previous work identified two populations of midline progenitors among the NMPs in zebrafish; a dorsal population that produces notochord and floor plate, and a ventral population that generates notochord and hypochord (Row et al., 2016). However, it remained unclear: 1) exactly where these progenitors reside in the tailbud, 2) how the progenitor cells move to generate these distinct midline cell types, and 3) how dorsal and ventral progenitors are maintained as distinct populations without mixing. For example, ventral midline progenitors never give rise to floor plate and dorsal midline progenitors never give rise to hypochord. Here, we pinpoint the locations of dorsal midline progenitors by first identifying cells with a high ratio of canonical Wnt signaling activity to *sox2* expression, similar to that of the posterior wall NMPs that generate spinal cord and paraxial/endothelial mesoderm. These dorsal midline NMP cells reside in the posterior-most region of what would have previously been described as the floor plate (Fig. 8A). Using midline targeted transplants and time-lapse imaging, we show that these cells migrate posteriorly with respect to notochord progenitors. Some of them slow their posterior migration and become floor plate, while others change shape and move ventrally to join the notochord (Fig. 8B). Using a Cdk activity sensor transgenic line, we also show that the dorsal and ventral midline progenitor populations proliferate and enter the G1 phase of the cell cycle before becoming notochord (Fig. 8C). scRNA-seq data provide additional support for this conclusion.

**Figure 8.**
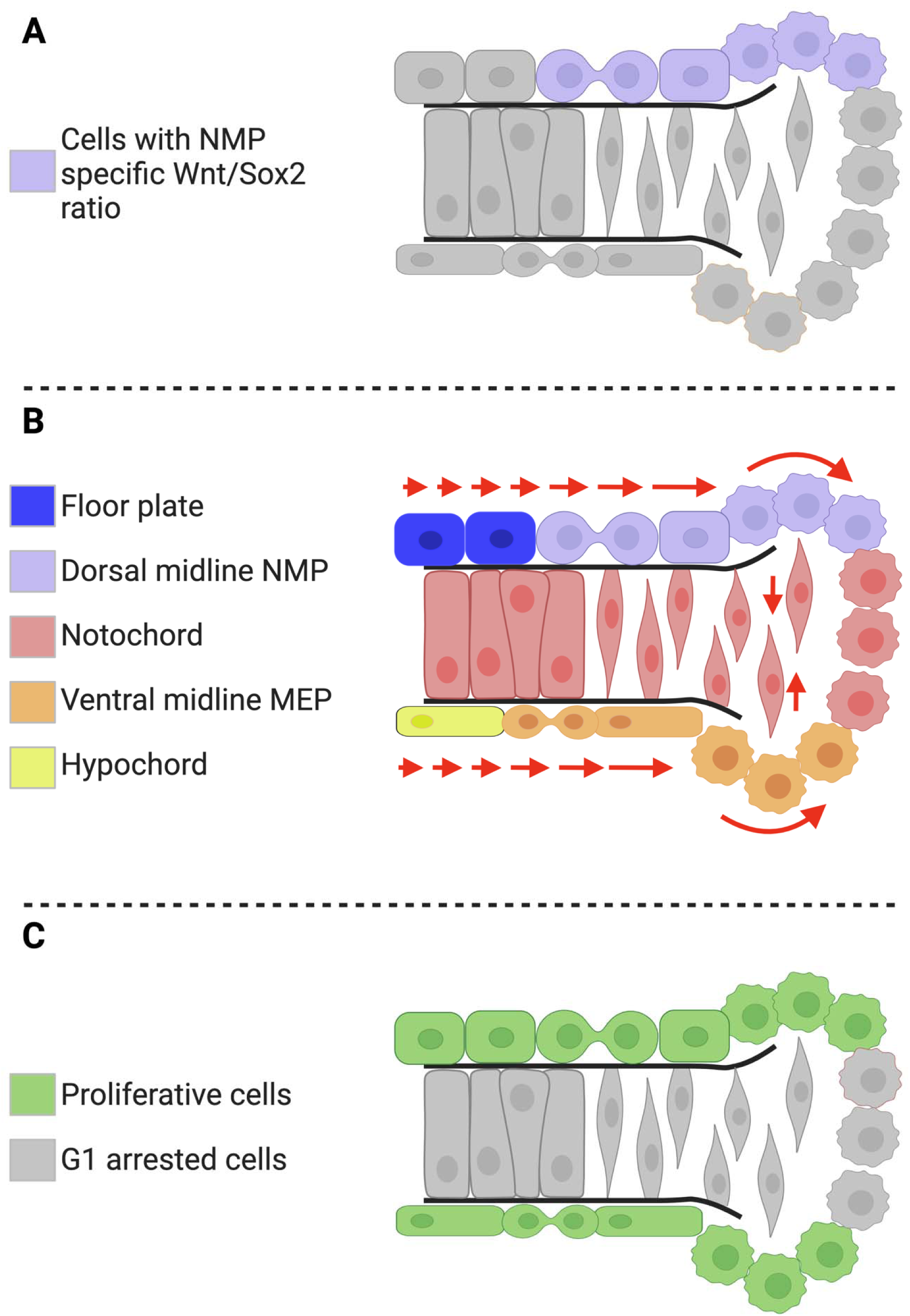
Model of zebrafish midline development. (A) The midline cells of a zebrafish embryo are schematized showing the location of cells (purple) with an NMP specific Wnt/Sox2 ratio. (B) These cells represent the dorsal midline NMPs (purple) that generate floor plate (blue) and notochord (red). Ventral mesendodermal progenitors (MEPs, orange) give rise to hypochord (yellow) and notochord (red). Red arrows show the movement of cells during tissue formation. NMPs and MEPs migrate posteriorly with some cells at the posterior-most domains converging and extending to form notochord, while cells in the trailing region slow their migration and form floor plate or hypochord, respectively. (C) Midline cells that are actively proliferating based on a Cdk activity reporter are highlighted in green.

In our experiments activating *sox2* expression mosaically, we observed cells accumulating at the position of the dorsal midline progenitors, as well as at the posterior-most hypochord, both regions of endogenous *sox2* expression. Time-lapse movies of transplanted cells show that the posterior hypochord region migrates posteriorly with respect to the notochord, and that cells from this region can slow their migration to generate hypochord or change shape and move dorsally to become notochord. Thus, our data suggest that the region previously thought to be the posterior-most hypochord contains the ventral midline progenitors. The physical separation and posterior-ward migration of the dorsal and ventral midline progenitors likely ensures that cells from the dorsal midline progenitor population cannot join the hypochord, and cells from the ventral midline progenitors cannot join the floor plate (Fig. 8B).

### Defining the neuromesodermal progenitor state based on the ratio of canonical Wnt signaling activity and *sox2* expression levels

One of the defining features of NMPs is their co-expression of the *sox2* and *tbxt* transcription factors. We previously showed that a primary function of *tbxt* is to activate the canonical Wnt signaling pathway through direct activation of Wnt ligand expression, including *wnt3a* and *wnt8a* (Martin and Kimelman, 2008). Wnt signaling in turn activates *tbxt* expression, creating an autoregulatory loop (Martin and Kimelman, 2008; Yamaguchi et al., 1999). In mosaic analyses in both mouse and zebrafish, posterior wall NMPs lacking Tbxt function are sustained over the course of axial development and can become mesoderm when surrounded by wild-type cells (Guibentif et al., 2021; Martin and Kimelman, 2008, 2010). However, when cells lack Wnt signaling they cannot form mesoderm, even when surrounded by wild-type cells (Martin and Kimelman, 2012). This suggests that Wnt signaling is a critical defining feature maintaining the NMP potential.

NMPs maintain *sox2* expression as they form neural tissue and rapidly downregulate it as they become mesoderm. Loss of *sox2* function in zebrafish or *Sox2* and *Sox3* in mouse NMPs disrupts somite development, indicating that SoxB1 genes play an important role in the maintenance of NMPs (Kinney et al., 2020; Yoshida et al., 2014). The transcription factors Tbx16 in zebrafish and TBX6 in mouse normally repress *sox2* transcription during the transition from NMP to mesoderm (Bouldin et al., 2015; Takemoto et al., 2011). Loss of function of these transcription factors prevents the majority of NMPs from transitioning to mesoderm and cells maintain their neuromesodermal potential (Chapman and Papaioannou, 1998; Griffin et al., 1998; Kimmel et al., 1989; Kinney et al., 2020; Takemoto et al., 2011). In zebrafish, when Wnt signaling is also inhibited in this context, NMPs exit into the anatomical position where mesoderm normally forms but instead these cells form ectopic neural tubes (Kinney et al., 2020), similar to the fate of *Tbx6* mutant cells in the mouse that manage to escape the tailbud into the region where mesoderm normally forms (Chapman and Papaioannou, 1998). These prior studies suggest that a unique interaction between Sox2 and canonical Wnt signaling restricts cells to the tailbud and maintains their neural and mesodermal fate potential. Here we show, using reporter transgenes, that NMPs in the posterior wall of the tailbud have a unique ratio of Wnt signaling activity to *sox2* expression level compared to newly induced neural or mesodermal tissue, and that this ratio can be used to predict and identify other cells with neuromesodermal potential. Thus, the co-activation of Wnt signaling and *sox2* expression can be considered a defining feature of NMPs. Since *Tbxt* is a direct Wnt target gene (Yamaguchi et al., 1999), it follows that NMPs also co-express *tbxt* and *sox2*.

Several studies have suggested models for how Wnt signaling and Sox2 may interact to generate a unique output in NMPs distinct from either Wnt or Sox2 alone. The Wnt effector protein β-catenin directly interacts with numerous Sox transcription factors, including Sox2, and this can influence gene expression patterns (Kormish et al., 2010; Ye et al., 2014). In pluripotent, stem cell-derived NMPs in vitro, the vast majority of genomic regions occupied by β-catenin are co-occupied by Sox2 (Mukherjee et al., 2022). When Sox2 levels are reduced, β-catenin binding is lost at many of these sites. Additionally, Sox2 levels can affect the occupancy of Sox2 at different genomic locations (Blassberg et al., 2022). This in turn changes the genomic occupancy of β-catenin, with β-catenin tending to co-occupy sites bound by Sox2. Thus, Sox2 can elicit unique responses to Wnt signaling when it is present, which can vary depending on the precise level of Sox2 and this likely establishes an NMP-specific transcriptional output.

### Auxin inducible protein degradation in zebrafish

Rapid conditional depletion of proteins is a powerful method for studying protein function. Transcription factors can be depleted in zebrafish using Transcription Factor Targeting Chimeras, but currently this system is not conditional (Samarasinghe et al., 2021). Methods to deplete GFP-tagged proteins in zebrafish include a GFP nanobody fused to the F-box protein Fbxw11b, which facilitates ubiquitination and subsequent degradation of GFP fusion proteins (Yamaguchi et al., 2019). An alternative nanobody approach utilized a GFP nanobody fused to an auxin inducible degron (AID) (Daniel et al., 2018). In the presence of the plant F-box protein Tir1, the AID-tagged nanobody and its bound GFP fusion protein are ubiquitinated and targeted for degradation following addition of the plant hormone auxin. A direct approach to tagging proteins of interest with AID, as opposed to a nanobody, has also been tested in zebrafish (Yamaguchi et al., 2019). However, degradation was leaky in the presence of Tir1 and absence of auxin. A direct AID tagging approach has been a powerful method of protein depletion in many systems including protozoans, mammalian cell culture, yeast, C. elegans, D. melanogaster, mouse, and others (Brown et al., 2017; Kanke et al., 2011; Kreidenweiss et al., 2013; Nishimura et al., 2009; Phanindhar and Mishra, 2023; Trost et al., 2016; Yesbolatova et al., 2020; Zhang et al., 2015). It provides rapid and reversible protein degradation based on the presence or absence of auxin. However, in some cases degradation with this system is also leaky and requires high concentrations of auxin. An updated system called AID 2 was developed using a bump and hole approach (Yesbolatova et al., 2020). A F74G mutation was introduced to the Tir1 protein, which binds strongly to the auxin derivative 5-Ph-IAA. The result is reduced leaky degradation and a requirement for 670-fold less ligand concentration.

We have established a modified direct AID tagging as a protein depletion method in zebrafish using many of the recent improvements made to the system, including the AID 2 approach (Yesbolatova et al., 2020). Instead of using Tir1 from *Oryza sativa*, which is commonly used in mammalian systems, we used the Tir1 from *Arabidopsis thaliana*, which is used in *C. elegans* (Zhang et al., 2015). *Arabidopsis thaliana* has a preferred temperature range of 23-25°C which is close to the preferred 28°C of zebrafish. We used a F79G mutant form of *A. thaliana* Tir1 (equivalent to the F74G mutation in *O. sativa* Tir1), paired with 5-Ph-IAA. We also introduced the D170E and M473L mutations, which were shown to improve binding of Tir1 to AID domains (Yu et al., 2013). Together this combination efficiently degrades tagged mNG fluorescent proteins. We also tested the mIAA7 degron, which was shown to provide a more rapid and complete degradation in mammalian cells and *C. elegans* (Li et al., 2019; Sepers et al., 2022), and this improved degradation kinetics. We expect the AID 2 system be broadly useful for the zebrafish community.

## Materials and Methods

### Zebrafish care and lines

All zebrafish methods were approved by the Stony Brook University Institutional Animal Care and Use Committee. Transgenic lines used include *Tg(hsp70l:sox2-2A-NLS-KikGR)*^sbu100^ (referred to here as *HS:sox2*) (Row et al., 2016), *Tg(hsp70l:Xla.Tcf.-EGFP)*^w74^ (referred to here as *HS:tcf*Δ*c*) (Martin and Kimelman, 2012), *Tg(sox2-2A-sfGFP)*^stl84^ (Shin et al., 2014), *Tg*(*7xTCF-Xla.Siam:nlsmCherry)* (Moro et al., 2012), *Tg(hsp70l:CAAX-mCherry-2A-NLS-KikGR)*^sbu104^ (referred to here as *HS:CAAX-mCherry-2A-NLS-KikGR*) (Goto et al., 2017), *tg(ubb:mmu.lck-mneongreen)*^sbu107^ (Adikes et al., 2020), *Tg(hsp70l:hsa.helb-mneongreen-2a-hsa.hist1h2bj-mscarlet)*^sbu108^ (referred to here as *HS:DHB-mNG-P2A-H2B-mSc*) (Adikes et al., 2020), and *Tg(hsp70l:hsa.helb-mscarlet-2a-hsa.hist1h2bj-mirfp670)*^sbu109^ (referred to here as *HS:DHB-mSc-2A-H2B-miRFP670*) (Adikes et al., 2020). All heat-shock inductions were performed by moving embryos from 28°C to prewarmed 40°C embryo media and held at 40°C for 30 minutes.

### Plasmid construction

The lab’s heat shock plasmid comprises the Hsp70 heat shock promoter positioned upstream of two specific restriction enzyme cut sites (BamHI and ClaI). These sites served the purpose of linearizing the heat shock vector for subsequent plasmid construction.

HS:Tir1-E2A-BFP (pRM30) construction: Hsp70 was digested with BamHI and ClaI to linearize the plasmid for assembly. Using NEB HiFi, the Tir1:E2A:BFP PCR product was inserted into the linearized Hsp70 plasmid to create the final HS:Tir1:E2A:BFP and TTCGTGGCTCCAGAGAATCGATattaagcttgtgccccagtttgcta. The forward primer contains homology to the Hsp70 BamHI cut site, and the reverse primer has homology to the ClaI cut site. Using NEB HiFi, the Tir1:E2A:BFP PCR product was inserted into the linearized Hsp70 plasmid to create the final HS:Tir1:E2A:BFP plasmid.

HS:Tir1-E2A-mScarlet (pRM31) construction:HS:Tir1-E2A-BFP was digested with MluI and ClaI to remove BFP and linearize the plasmid for assembly. The E2A:mScarlet sequence was amplified from pRM25 plasmid using primers (RM485/486): TCATCACCACCAACGGACTCACGCGTGGCAGCGGCCAGTG and CTTATCATGTCTGGATCATCATCGATTTACTTGTACAGCTCGTCCA. The forward primer contains homology to the Hsp70 MluI cut site, and the reverse primer has homology to the ClaI cut site. Using NEB HiFi, the E2A:mScarlet PCR product was inserted into the linearized Hsp70-aid plasmid to create the final HS:Tir1:E2A:mScarlet plasmid.

HS:mNG-linker-aid (pRM33) construction: The Hsp70 plasmid was digested with BamHI and ClaI enzymes to linearize it for subsequent assembly. The mNG:linker:aid sequence was then amplified from a previously constructed plasmid using the primers CAAGCTACTTGTTCTTTTTGCAGGATCCGCCGCCACCATGGTGAGCAAGGGCGA and TCGTGGCTCCAGAGAATCGATttaTTTGACAAACGCGGCTGC (RM516/RM517). The forward primer was designed with homology to the BamHI cut site within Hsp70, while the reverse primer was designed with homology to the ClaI cut site. Utilizing NEB Hifi, the resulting mNG-linker-aid PCR product was inserted into the linearized Hsp70 plasmid, yielding the final construct termed HSLmNG-linker-aid.

HS:mNG-linker-Sox2-linker-aid (pRM34) construction:Hsp70 plasmid was digested with BamHI and ClaI enzymes to linearize it for subsequent assembly. The mNG sequence was then amplified using the primers CAAGCTACTTGTTCTTTTTGCAGGATCCGCCGCCACCATGGTGAGCAAGGGCG and TtgcaccGCTAGCaccACCGGTCTTGTACAGCTCGTCCATG (RM437/441), the forward primer added homology to the vector and the reverse primer added homology to the linker upstream of Sox2. Sox2 was amplified using the primers ggtGCTAGCggtgcaAGCggcgccAGCATGGATTATAAGGATCACGATGGAG and TGTGGCTGAGTGGTATGGTT (RM470/471), with the forward primer adding a 5’ linker to the Sox2 PCR product and the reverse primer having no homology. PCR was performed on the Sox2 PCR product to add homology for subsequent assembly. The mIAA7aid sequence was amplified using the forward primer GACGAGCTGTACAAGtaaACCGGTggtGCTAGCggtgcaAGC and the reverse primer TCGTGGCTCCAGAGA ATCGAT TTTTGACAAACGCGGCTGCC (RM493/RM440). Utilizing NEB Hifi polymerase, all three PCR products were inserted into the linearized Hsp70 plasmid to create the final construct, termed HS:mNG-linker-Sox2-linker-aid vector. The assembly of all three PCR products into the Hsp70 BamHI/Clal cut site was achieved using NEB Hifi assembly method.

HS:mNG-linker-Sox2-linker-mIAA7aid-H2B-mScarlet (pRM35) construction: HS:mNG-linker-Sox2-linker-aid was digested with ClaI to linearize the plasmid for subsequent assembly. The P2A:H2B:mScarlet sequence was then amplified using the primers GACCCAACAAAAGACCTCCTCCATCGATGGCAGCGGCGCCACC and GAGAAGTTCGTGGCTCCAGAGAATCGATttaCTTGTACAGCTCGTCCATGCCG (RM510/511). The forward primer was designed with homology to the sequence upstream of the ClaI cut site in HS:mNG-linker-Sox2-linker-aid, while the reverse primer contained homology downstream of the ClaI cut site in HS:mNG-linker-Sox2-linker-aid.Utilizing NEB Hifi, the resulting P2A-H2B-mScarlet PCR product was inserted into the linearized plasmid, resulting in the creation of the final construct, HS:mNG-linker-Sox2-linker-mIAA7aid.

### Generation of the *HS:tir1-E2A-BFP* transgenic line

The stable zebrafish line *HS:tir1-E2A-BFP* was generated utilizing the Tol2 recombinase system (Kawakami, 2004). The lines were established by injecting single-cell wild-type zebrafish embryos with 25 pg of *HS:tir1-E2A-BFP* plasmid along with 25 pg of *tol2* mRNA, employing previously described methods. F0 injected embryos were subsequently crossed with wild-type zebrafish, screening for germline transmission of the transgene.

### 5-Ph-IAA (Auxin) treatment

In all auxin experiments, 5-Ph-IAA auxin was resuspended in 100% DMSO at a concentration of 100 mM and stored at −20°C. Zebrafish embryos were treated by directly adding auxin to embryo media at a concentration of 1 mM. Additionally, when necessary, 5-Ph-IAA at a concentration of 2 mM was added to 1% low melting point agar, prior to imaging. During 5-Ph-IAA treatments, 100% DMSO was added to increase tissue permeabilization at a final concentration of 2%. Furthermore, immobilization of embryos for imaging was achieved using a concentration of 1x tricaine (25x stock; 0.4 g/l; Pentair, TRS1). For time-lapse imaging, embryos were treated during mounting, immediately preceding imaging while end-point experiments embryos were treated at the bud stage.

### Image quantification

Image quantification was manually carried out using Fiji (Schindelin et al., 2012). To address the significant amplifier noise in EM-CCD images and eliminate residual out-of-focus fluorescence in these confocal micrographs, background subtraction was performed. The Z plane containing the cell’s center of interest was identified. The freehand tool was used to draw an area for measurement. The mean gray value within the region was measured. This procedure was repeated for each cell of interest in the image. For time-lapse recordings, the procedure was repeated at each time point following individual cells.

### Embryo mounting and microscopy

Prior to imaging, live zebrafish embryos were immobilized using a solution of 1% low melting point agarose dissolved in embryo media supplemented with 1x tricaine (25x stock 0.4 g/l; Pentair, TRS1) and 2% DMSO. Static tailbud fluorescent images were captured between 13 and 17 somites using a 40x magnification dip lens with a Hamamatsu Orca EM-CCD camera and a Borealis-modified Yokagawa CSU-10 spinning disk confocal microscope (Nobska Imaging, Inc.). Time-lapse movies of individual cells were recorded starting at either 6 or 12 somites using the same imaging setup. Subsequent image processing was performed using Fiji.

### Hybridization chain reaction

Version 3 HCR was performed using the protocol called “HCR™ RNA-FISH protocol for whole-mount zebrafish embryos and larvae (*Danio rerio*)” provided by Molecular Instruments and available for download on their website (Choi et al., 2018). The only modification we made was we used 5ul of stock hairpin during the amplification stage whereas the protocol calls for 10ul.

### Single-cell RNA sequencing

Wild type AB zebrafish embryos were obtained by natural breeding. Tailbuds were obtained by manual dissection at 16 hpf. Tailbuds were then dissociated into single cell suspension using mechanical disruption and trypsin/collagenase P incubation as described in Barske et al. (Barske et al., 2016). Single cell suspensions were processed on a 10x Chromium platform for single-cell library construction. Libraries were then sequenced on a HiSeq2500 (Illumina). FASTQ files were mapped to the zebrafish transcriptome GRCz11 using CellRanger. Mapped reads for the 12-32 hpf timeline were aggregated using CellRanger before further computational analysis. Counts matrix normalization and scaling were performed using Seurat v3.

### Computational analysis and pseudotime computation of single-cell RNA-seq data

Variable features identification, principal component analysis (PCA), UMAP reduction, and unbiased clustering were performed in Seurat v3 (Butler et al., 2018). Differential expression analysis in Seurat v3 was used to identify cluster markers by wilcoxon rank sum test. Cluster markers were then used to assign cell types. Visualizations were generated using Seurat v3 and ggplot2 with color palettes from the RColorBrewer and Viridis packages. A subset counts matrix containing TFs was analyzed with Monocle3 for trajectory and pseudotime analysis (Trapnell et al., 2014). Identification of temporally distinct TF modules was performed using the graph_test and find_gene_modules functions in Monocle3.

## Acknowledgements

We thank Stephanie Flanagan and Calvin Yu for zebrafish care, David Matus for discussions and advice about the auxin inducible degron system, and group members for critical feedback. All figures were generated with Biorender. This work was supported by grants from the Royal Society of New Zealand (MFP-UOO2013) and the Fulbright NZ to JAH, from the NIDCR (R01DE13828, R01DE30565), NSF (MCB2028424), and Simons Foundation (594598) to TFS, from the NICHD (F31HD108921) to SS, and from the NIGMS (R01GM124282 and R35GM150290) to BLM.

## References

Adikes, R.C., Kohrman, A.Q., Martinez, M.A.Q., Palmisano, N.J., Smith, J.J., Medwig-Kinney, T.N., Min, M., Sallee, M.D., Ahmed, O.B., Kim, N., Liu, S., Morabito, R.D., Weeks, N., Zhao, Q., Zhang, W., Feldman, J.L., Barkoulas, M., Pani, A.M., Spencer, S.L., Martin, B.L., Matus, D.Q., 2020. Visualizing the metazoan proliferation-quiescence decision in vivo. Elife 9 (10.7554/eLife.63265).

Amacher, S.L., Kimmel, C.B., 1998. Promoting notochord fate and repressing muscle development in zebrafish axial mesoderm. Development 125, 1397–1406.

Avilion, A.A., Nicolis, S.K., Pevny, L.H., Perez, L., Vivian, N., Lovell-Badge, R., 2003. Multipotent cell lineages in early mouse development depend on SOX2 function. Genes Dev 17, 126–140 (10.1101/gad.224503).

Barske, L., Askary, A., Zuniga, E., Balczerski, B., Bump, P., Nichols, J.T., Crump, J.G., 2016. Competition between Jagged-Notch and Endothelin1 Signaling Selectively Restricts Cartilage Formation in the Zebrafish Upper Face. PLoS Genet 12, e1005967 (10.1371/journal.pgen.1005967).

Bergsland, M., Ramskold, D., Zaouter, C., Klum, S., Sandberg, R., Muhr, J., 2011. Sequentially acting Sox transcription factors in neural lineage development. Genes Dev 25, 2453–2464 (10.1101/gad.176008.111).

Blassberg, R., Patel, H., Watson, T., Gouti, M., Metzis, V., Delas, M.J., Briscoe, J., 2022. Sox2 levels regulate the chromatin occupancy of WNT mediators in epiblast progenitors responsible for vertebrate body formation. Nat Cell Biol 24, 633–644 (10.1038/s41556-022-00910-2).

Bouldin, C.M., Manning, A.J., Peng, Y.H., Farr, G.H., Hung, K.L., Dong, A., Kimelman, D., 2015. Wnt signaling and tbx16 form a bistable switch to commit bipotential progenitors to mesoderm. Development 142, 2499-+ (10.1242/dev.124024).

Brown, K.M., Long, S., Sibley, L.D., 2017. Plasma Membrane Association by N-Acylation Governs PKG Function in Toxoplasma gondii. mBio 8 (10.1128/mBio.00375-17).

Bunina, D., Abazova, N., Diaz, N., Noh, K.M., Krijgsveld, J., Zaugg, J.B., 2020. Genomic Rewiring of SOX2 Chromatin Interaction Network during Differentiation of ESCs to Postmitotic Neurons. Cell Syst 10, 480–494 e488 (10.1016/j.cels.2020.05.003).

Butler, A., Hoffman, P., Smibert, P., Papalexi, E., Satija, R., 2018. Integrating single-cell transcriptomic data across different conditions, technologies, and species. Nature biotechnology 36, 411–420 (10.1038/nbt.4096).

Catala, M., Teillet, M.A., Le Douarin, N.M., 1995. Organization and development of the tail bud analyzed with the quail-chick chimaera system. Mech Dev 51, 51–65.

Chapman, D.L., Papaioannou, V.E., 1998. Three neural tubes in mouse embryos with mutations in the T-box gene Tbx6. Nature 391, 695–697.

Choi, H.M.T., Schwarzkopf, M., Fornace, M.E., Acharya, A., Artavanis, G., Stegmaier, J., Cunha, A., Pierce, N.A., 2018. Third-generation in situ hybridization chain reaction: multiplexed, quantitative, sensitive, versatile, robust. Development 145 (10.1242/dev.165753).

Daniel, K., Icha, J., Horenburg, C., Muller, D., Norden, C., Mansfeld, J., 2018. Conditional control of fluorescent protein degradation by an auxin-dependent nanobody. Nat Commun 9, 3297 (10.1038/s41467-018-05855-5).

Ekker, S.C., Ungar, A.R., Greenstein, P., von Kessler, D.P., Porter, J.A., Moon, R.T., Beachy, P.A., 1995. Patterning activities of vertebrate hedgehog proteins in the developing eye and brain. Curr Biol 5, 944–955 (10.1016/s0960-9822(95)00185-0).

Genuth, M.A., Kojima, Y., Julich, D., Kiryu, H., Holley, S.A., 2023. Automated time-lapse data segmentation reveals in vivo cell state dynamics. Sci Adv 9, eadf1814 (10.1126/sciadv.adf1814).

Goto, H., Kimmey, S.C., Row, R.H., Matus, D.Q., Martin, B.L., 2017. FGF and canonical Wnt signaling cooperate to induce paraxial mesoderm from tailbud neuromesodermal progenitors through regulation of a two-step epithelial to mesenchymal transition. Development 144, 1412–1424 (10.1242/dev.143578).

Griffin, K.J., Amacher, S.L., Kimmel, C.B., Kimelman, D., 1998. Molecular identification of spadetail: regulation of zebrafish trunk and tail mesoderm formation by T-box genes. Development 125, 3379–3388.

Guibentif, C., Griffiths, J.A., Imaz-Rosshandler, I., Ghazanfar, S., Nichols, J., Wilson, V., Gottgens, B., Marioni, J.C., 2021. Diverse Routes toward Early Somites in the Mouse Embryo. Dev Cell 56, 141–153 e146 (10.1016/j.devcel.2020.11.013).

Halpern, M.E., Thisse, C., Ho, R.K., Thisse, B., Riggleman, B., Trevarrow, B., Weinberg, E.S., Postlethwait, J.H., Kimmel, C.B., 1995. Cell-autonomous shift from axial to paraxial mesodermal development in zebrafish floating head mutants. Development 121, 4257–4264.

Kanke, M., Nishimura, K., Kanemaki, M., Kakimoto, T., Takahashi, T.S., Nakagawa, T., Masukata, H., 2011. Auxin-inducible protein depletion system in fission yeast. BMC Cell Biol 12, 8 (10.1186/1471-2121-12-8).

Kawakami, K., 2004. Transgenesis and gene trap methods in zebrafish by using the Tol2 transposable element. Methods Cell Biol 77, 201–222.

Kimelman, D., 2016. Tales of Tails (and Trunks): Forming the Posterior Body in Vertebrate Embryos. Curr Top Dev Biol 116, 517–536 (10.1016/bs.ctdb.2015.12.008).

Kimelman, D., Martin, B.L., 2012. Anterior-posterior patterning in early development: three strategies. Wiley Interdiscip Rev Dev Biol 1, 253–266 (10.1002/wdev.25).

Kimmel, C.B., Kane, D.A., Walker, C., Warga, R.M., Rothman, M.B., 1989. A mutation that changes cell movement and cell fate in the zebrafish embryo. Nature 337, 358–362 (10.1038/337358a0).

Kinney, B.A., Al Anber, A., Row, R.H., Tseng, Y.J., Weidmann, M.D., Knaut, H., Martin, B.L., 2020. Sox2 and Canonical Wnt Signaling Interact to Activate a Developmental Checkpoint Coordinating Morphogenesis with Mesoderm Fate Acquisition. Cell Rep 33, 108311 (10.1016/j.celrep.2020.108311).

Koch, F., Scholze, M., Wittler, L., Schifferl, D., Sudheer, S., Grote, P., Timmermann, B., Macura, K., Herrmann, B.G., 2017. Antagonistic Activities of Sox2 and Brachyury Control the Fate Choice of Neuro-Mesodermal Progenitors. Dev Cell 42, 514–526 e517 (10.1016/j.devcel.2017.07.021).

Kormish, J.D., Sinner, D., Zorn, A.M., 2010. Interactions between SOX factors and Wnt/beta-catenin signaling in development and disease. Dev Dyn 239, 56–68 (10.1002/dvdy.22046).

Kreidenweiss, A., Hopkins, A.V., Mordmuller, B., 2013. 2A and the auxin-based degron system facilitate control of protein levels in Plasmodium falciparum. PLoS One 8, e78661 (10.1371/journal.pone.0078661).

La Manno, G., Soldatov, R., Zeisel, A., Braun, E., Hochgerner, H., Petukhov, V., Lidschreiber, K., Kastriti, M.E., Lonnerberg, P., Furlan, A., Fan, J., Borm, L.E., Liu, Z., van Bruggen, D., Guo, J., He, X., Barker, R., Sundstrom, E., Castelo-Branco, G., Cramer, P., Adameyko, I., Linnarsson, S., Kharchenko, P.V., 2018. RNA velocity of single cells. Nature 560, 494–498 (10.1038/s41586-018-0414-6).

Lange, M., Granados, A., VijayKumar, S., Bragantini, J., Ancheta, S., Kim, Y.J., Santhosh, S., Borja, M., Kobayashi, H., McGeever, E., Solak, A.C., Yang, B., Zhao, X., Liu, Y., Detweiler, A.M., Paul, S., Theodoro, I., Mekonen, H., Charlton, C., Lao, T., Banks, R., Xiao, S., Jacobo, A., Balla, K., Awayan, K., D’Souza, S., Haase, R., Dizeux, A., Pourquie, O., Gomez-Sjoberg, R., Huber, G., Serra, M., Neff, N., Pisco, A.O., Royer, L.A., 2024. A multimodal zebrafish developmental atlas reveals the state-transition dynamics of late-vertebrate pluripotent axial progenitors. Cell 187, 6742–6759 e6717 (10.1016/j.cell.2024.09.047).

Le Douarin, N.M., Halpern, M.E., 2000. Discussion point. Origin and specification of the neural tube floor plate: insights from the chick and zebrafish. Current opinion in neurobiology 10, 23–30.

Li, S., Prasanna, X., Salo, V.T., Vattulainen, I., Ikonen, E., 2019. An efficient auxin-inducible degron system with low basal degradation in human cells. Nat Methods 16, 866–869 (10.1038/s41592-019-0512-x).

Martin, B., 2020. Progenitor Cells in Vertebrate Segmentation, in: Chipman, A. (Ed.), Cellular Processes in Segmentation, 1st ed. CRC Press, Boca Raton, pp. 99–123.

Martin, B.L., 2021. Mesoderm induction and patterning: Insights from neuromesodermal progenitors. Semin Cell Dev Biol (10.1016/j.semcdb.2021.11.010).

Martin, B.L., Kimelman, D., 2008. Regulation of canonical Wnt signaling by Brachyury is essential for posterior mesoderm formation. Dev Cell 15, 121–133 (10.1016/j.devcel.2008.04.013).

Martin, B.L., Kimelman, D., 2009. Wnt signaling and the evolution of embryonic posterior development. Curr Biol 19, R215–219 (10.1016/j.cub.2009.01.052).

Martin, B.L., Kimelman, D., 2010. Brachyury establishes the embryonic mesodermal progenitor niche. Genes Dev 24, 2778–2783 (10.1101/gad.1962910).

Martin, B.L., Kimelman, D., 2012. Canonical Wnt signaling dynamically controls multiple stem cell fate decisions during vertebrate body formation. Dev Cell 22, 223–232 (10.1016/j.devcel.2011.11.001).

Martin, B.L., Steventon, B., 2022. A fishy tail: Insights into the cell and molecular biology of neuromesodermal cells from zebrafish embryos. Dev Biol 487, 67–73 (10.1016/j.ydbio.2022.04.010).

Morabito, R.D., Adikes, R.C., Matus, D.Q., Martin, B.L., 2021. Cyclin-Dependent Kinase Sensor Transgenic Zebrafish Lines for Improved Cell Cycle State Visualization in Live Animals. Zebrafish (10.1089/zeb.2021.0059).

Moro, E., Ozhan-Kizil, G., Mongera, A., Beis, D., Wierzbicki, C., Young, R.M., Bournele, D., Domenichini, A., Valdivia, L.E., Lum, L., Chen, C., Amatruda, J.F., Tiso, N., Weidinger, G., Argenton, F., 2012. In vivo Wnt signaling tracing through a transgenic biosensor fish reveals novel activity domains. Developmental biology 366, 327–340 (10.1016/j.ydbio.2012.03.023).

Mukherjee, S., Luedeke, D.M., McCoy, L., Iwafuchi, M., Zorn, A.M., 2022. SOX transcription factors direct TCF-independent WNT/beta-catenin responsive transcription to govern cell fate in human pluripotent stem cells. Cell Rep 40, 111247 (10.1016/j.celrep.2022.111247).

Nishimura, K., Fukagawa, T., Takisawa, H., Kakimoto, T., Kanemaki, M., 2009. An auxin-based degron system for the rapid depletion of proteins in nonplant cells. Nat Methods 6, 917–922 (10.1038/nmeth.1401).

Phanindhar, K., Mishra, R.K., 2023. Auxin-inducible degron system: an efficient protein degradation tool to study protein function. Biotechniques 74, 186–198 (10.2144/btn-2022-0108).

Row, R.H., Tsotras, S.R., Goto, H., Martin, B.L., 2016. The zebrafish tailbud contains two independent populations of midline progenitor cells that maintain long-term germ layer plasticity and differentiate in response to local signaling cues. Development 143, 244–254 (10.1242/dev.129015).

Samarasinghe, K.T.G., Jaime-Figueroa, S., Burgess, M., Nalawansha, D.A., Dai, K., Hu, Z., Bebenek, A., Holley, S.A., Crews, C.M., 2021. Targeted degradation of transcription factors by TRAFTACs: TRAnscription Factor TArgeting Chimeras. Cell Chem Biol 28, 648–661 e645 (10.1016/j.chembiol.2021.03.011).

Schindelin, J., Arganda-Carreras, I., Frise, E., Kaynig, V., Longair, M., Pietzsch, T., Preibisch, S., Rueden, C., Saalfeld, S., Schmid, B., Tinevez, J.Y., White, D.J., Hartenstein, V., Eliceiri, K., Tomancak, P., Cardona, A., 2012. Fiji: an open-source platform for biological-image analysis. Nat Methods 9, 676–682 (10.1038/nmeth.2019).

Sepers, J.J., Verstappen, N.H.M., Vo, A.A., Ragle, J.M., Ruijtenberg, S., Ward, J.D., Boxem, M., 2022. The mIAA7 degron improves auxin-mediated degradation in Caenorhabditiselegans. G3 (Bethesda) 12 (10.1093/g3journal/jkac222).

Shin, J., Chen, J., Solnica-Krezel, L., 2014. Efficient homologous recombination-mediated genome engineering in zebrafish using TALE nucleases. Development 141, 3807–3818 (10.1242/dev.108019).

Spencer, S.L., Cappell, S.D., Tsai, F.C., Overton, K.W., Wang, C.L., Meyer, T., 2013. The proliferation-quiescence decision is controlled by a bifurcation in CDK2 activity at mitotic exit. Cell 155, 369–383 (10.1016/j.cell.2013.08.062).

Sugiyama, M., Saitou, T., Kurokawa, H., Sakaue-Sawano, A., Imamura, T., Miyawaki, A., Iimura, T., 2014. Live imaging-based model selection reveals periodic regulation of the stochastic G1/S phase transition in vertebrate axial development. PLoS Comput Biol 10, e1003957 (10.1371/journal.pcbi.1003957).

Sugiyama, M., Sakaue-Sawano, A., Iimura, T., Fukami, K., Kitaguchi, T., Kawakami, K., Okamoto, H., Higashijima, S., Miyawaki, A., 2009. Illuminating cell-cycle progression in the developing zebrafish embryo. Proc Natl Acad Sci U S A 106, 20812–20817 (10.1073/pnas.0906464106).

Takahashi, K., Tanabe, K., Ohnuki, M., Narita, M., Ichisaka, T., Tomoda, K., Yamanaka, S., 2007. Induction of pluripotent stem cells from adult human fibroblasts by defined factors. Cell 131, 861–872 (10.1016/j.cell.2007.11.019).

Takahashi, K., Yamanaka, S., 2006. Induction of pluripotent stem cells from mouse embryonic and adult fibroblast cultures by defined factors. Cell 126, 663–676 (10.1016/j.cell.2006.07.024).

Takemoto, T., Uchikawa, M., Yoshida, M., Bell, D.M., Lovell-Badge, R., Papaioannou, V.E., Kondoh, H., 2011. Tbx6-dependent Sox2 regulation determines neural or mesodermal fate in axial stem cells. Nature 470, 394–398 (10.1038/nature09729).

Talbot, W.S., Trevarrow, B., Halpern, M.E., Melby, A.E., Farr, G., Postlethwait, J.H., Jowett, T., Kimmel, C.B., Kimelman, D., 1995. A homeobox gene essential for zebrafish notochord development. Nature 378, 150–157 (10.1038/378150a0).

Teillet, M.A., Lapointe, F., Le Douarin, N.M., 1998. The relationships between notochord and floor plate in vertebrate development revisited. Proc Natl Acad Sci U S A 95, 11733–11738.

Thomson, M., Liu, S.J., Zou, L.N., Smith, Z., Meissner, A., Ramanathan, S., 2011. Pluripotency factors in embryonic stem cells regulate differentiation into germ layers. Cell 145, 875–889 (10.1016/j.cell.2011.05.017).

Trapnell, C., Cacchiarelli, D., Grimsby, J., Pokharel, P., Li, S., Morse, M., Lennon, N.J., Livak, K.J., Mikkelsen, T.S., Rinn, J.L., 2014. The dynamics and regulators of cell fate decisions are revealed by pseudotemporal ordering of single cells. Nature biotechnology 32, 381–386 (10.1038/nbt.2859).

Trost, M., Blattner, A.C., Lehner, C.F., 2016. Regulated protein depletion by the auxin-inducible degradation system in Drosophila melanogaster. Fly (Austin) 10, 35–46 (10.1080/19336934.2016.1168552).

Wang, Z., Oron, E., Nelson, B., Razis, S., Ivanova, N., 2012. Distinct lineage specification roles for NANOG, OCT4, and SOX2 in human embryonic stem cells. Cell Stem Cell 10, 440–454 (10.1016/j.stem.2012.02.016).

Wymeersch, F.J., Wilson, V., Tsakiridis, A., 2021. Understanding axial progenitor biology in vivo and in vitro. Development 148 (10.1242/dev.180612).

Yamaguchi, N., Colak-Champollion, T., Knaut, H., 2019. zGrad is a nanobody-based degron system that inactivates proteins in zebrafish. Elife 8 (10.7554/eLife.43125).

Yamaguchi, T.P., Takada, S., Yoshikawa, Y., Wu, N.Y., McMahon, A.P., 1999. T (Brachyury) is a direct target of Wnt3a during paraxial mesoderm specification. Genes & Development 13, 3185–3190 (10.1101/gad.13.24.3185).

Yan, Y.L., Hatta, K., Riggleman, B., Postlethwait, J.H., 1995. Expression of a type II collagen gene in the zebrafish embryonic axis. Dev Dyn 203, 363–376 (10.1002/aja.1002030308).

Ye, X., Wu, F., Wu, C., Wang, P., Jung, K., Gopal, K., Ma, Y., Li, L., Lai, R., 2014. beta-Catenin, a Sox2 binding partner, regulates the DNA binding and transcriptional activity of Sox2 in breast cancer cells. Cell Signal 26, 492–501 (10.1016/j.cellsig.2013.11.023).

Yesbolatova, A., Saito, Y., Kitamoto, N., Makino-Itou, H., Ajima, R., Nakano, R., Nakaoka, H., Fukui, K., Gamo, K., Tominari, Y., Takeuchi, H., Saga, Y., Hayashi, K.I., Kanemaki, M.T., 2020. The auxin-inducible degron 2 technology provides sharp degradation control in yeast, mammalian cells, and mice. Nat Commun 11, 5701 (10.1038/s41467-020-19532-z).

Yoshida, M., Uchikawa, M., Rizzoti, K., Lovell-Badge, R., Takemoto, T., Kondoh, H., 2014. Regulation of mesodermal precursor production by low-level expression of B1 Sox genes in the caudal lateral epiblast. Mech Dev 132, 59–68 (10.1016/j.mod.2014.01.003).

Yu, H., Moss, B.L., Jang, S.S., Prigge, M., Klavins, E., Nemhauser, J.L., Estelle, M., 2013. Mutations in the TIR1 auxin receptor that increase affinity for auxin/indole-3-acetic acid proteins result in auxin hypersensitivity. Plant Physiol 162, 295–303 (10.1104/pp.113.215582).

Zhang, L., Ward, J.D., Cheng, Z., Dernburg, A.F., 2015. The auxin-inducible degradation (AID) system enables versatile conditional protein depletion in C. elegans. Development 142, 4374–4384 (10.1242/dev.129635).

Zhang, S., Bell, E., Zhi, H., Brown, S., Imran, S.A.M., Azuara, V., Cui, W., 2019. OCT4 and PAX6 determine the dual function of SOX2 in human ESCs as a key pluripotent or neural factor. Stem Cell Res Ther 10, 122 (10.1186/s13287-019-1228-7).

Zhang, S., Cui, W., 2014. Sox2, a key factor in the regulation of pluripotency and neural differentiation. World J Stem Cells 6, 305–311 (10.4252/wjsc.v6.i3.305).

