## Supplementary figures and images for "The ratio of Wnt signaling activity to Sox2 transcription factor levels predicts neuromesodermal fate potential"

### Fig. S1

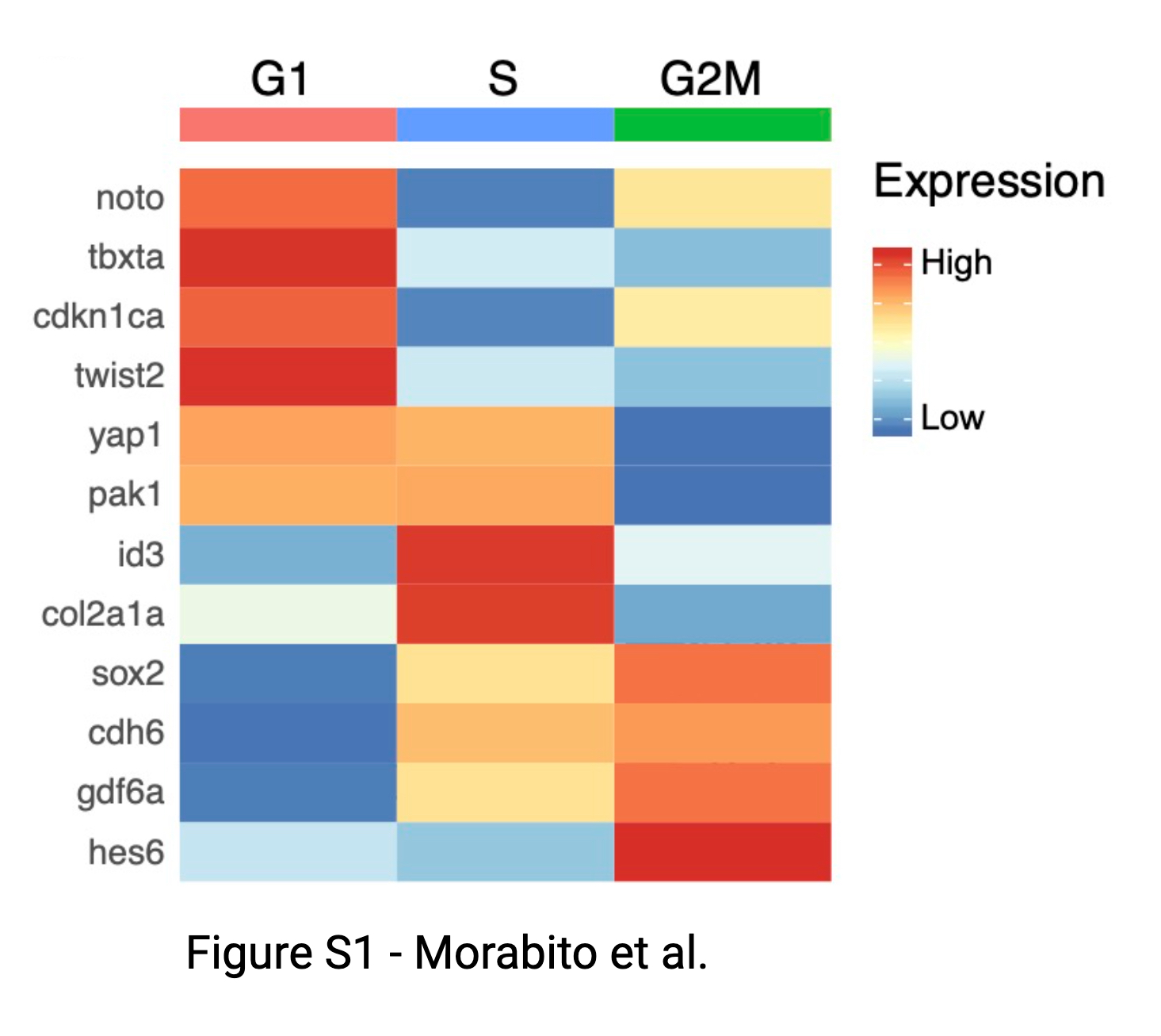

### Fig. S2

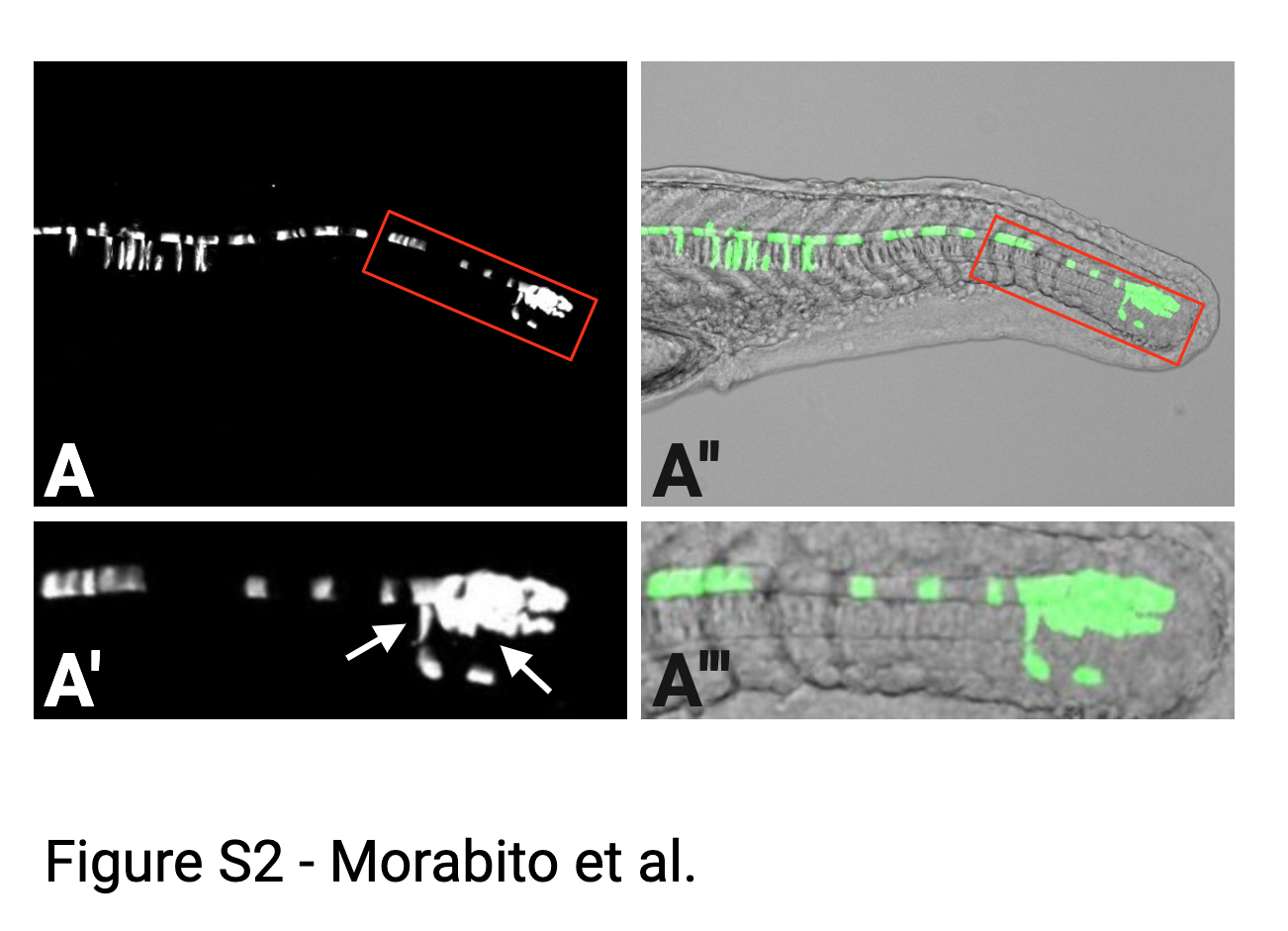

### Fig. S3

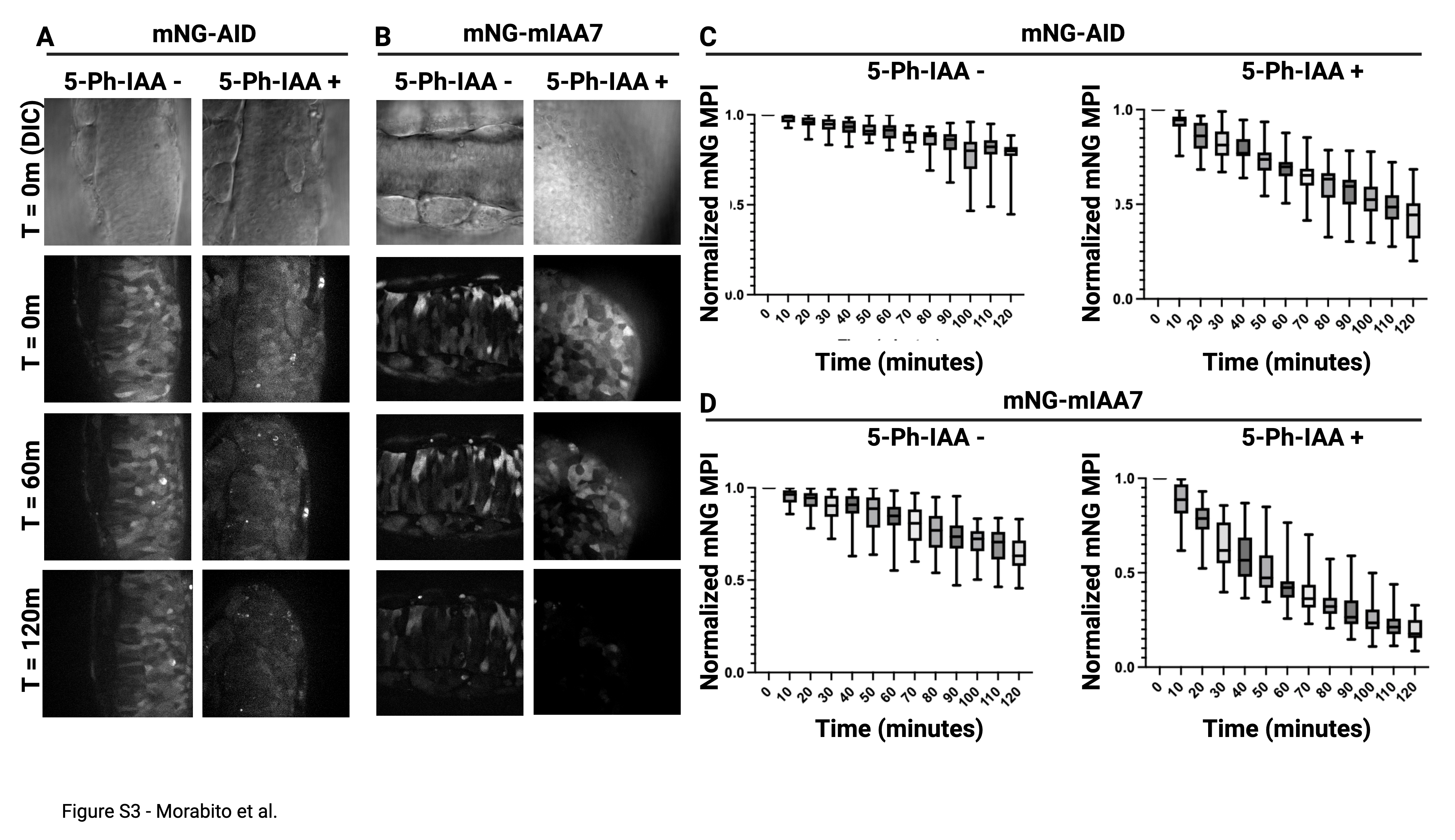

### Fig. S4

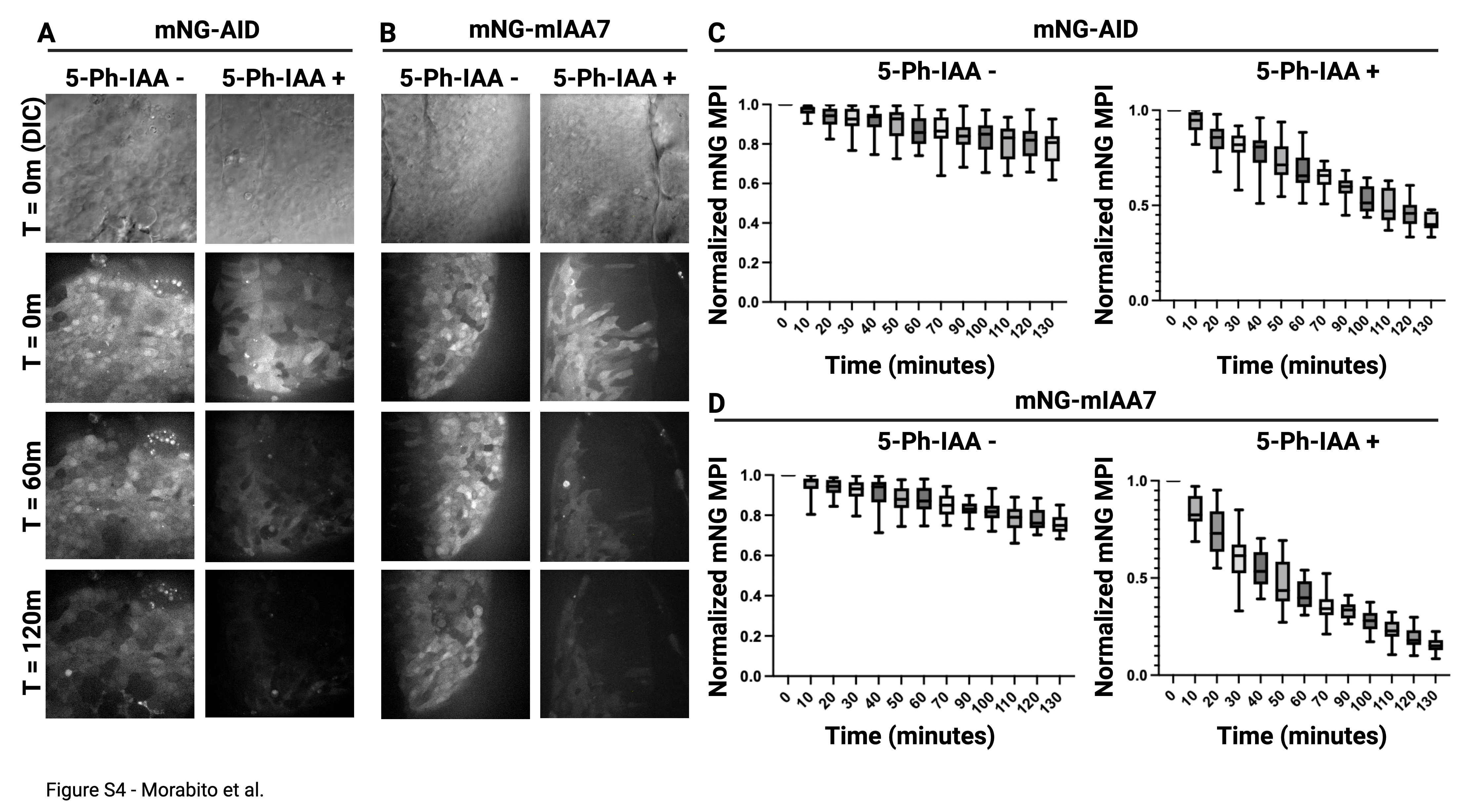
